# Cellular cartography reveals mouse prostate organization and determinants of castration resistance

**DOI:** 10.1101/2024.12.27.630532

**Authors:** Hanbyul Cho, Yuping Zhang, Jean C. Tien, Rahul Mannan, Jie Luo, Sathiya Pandi Narayanan, Somnath Mahapatra, Jing Hu, Greg Shelley, Gabriel Cruz, Miriam Shahine, Lisha Wang, Fengyun Su, Rui Wang, Xuhong Cao, Saravana Mohan Dhanasekaran, Evan T. Keller, Sethuramasundaram Pitchiaya, Arul M. Chinnaiyan

**Affiliations:** Michigan Center for Translational Pathology, University of Michigan, Ann Arbor, MI, 48109; Department of Pathology, University of Michigan, Ann Arbor, MI, 48109; Department of Computational Medicine and Bioinformatics, University of Michigan, Ann Arbor, MI 48109; Department of Pathology, Qilu Hospital, Cheeloo College of Medicine, Shandong University, Jinan, China; Department of Urology, University of Michigan, Ann Arbor, MI, 48109; Rogel Cancer Center, University of Michigan, Ann Arbor, MI, 48109; Single Cell Spatial Analysis Program, University of Michigan, Ann Arbor, MI, 48109; Biointerfaces Institute, University of Michigan, Ann Arbor, MI, 48109; Howard Hughes Medical Institute, University of Michigan, Ann Arbor, MI, 48109

**Keywords:** Single-cell RNA Sequencing, Single-cell ATAC Sequencing, Single-cell Multiomics, Spatial Transcriptomics, Androgen Signalling, Stress Response, Stemness, Castration Resistant Prostate cancer

## Abstract

Inadequate response to androgen deprivation therapy (ADT) frequently arises in prostate cancer, driven by cellular mechanisms that remain poorly understood. Here, we integrated single-cell RNA sequencing, single-cell multiomics, and spatial transcriptomics to define the transcriptional, epigenetic, and spatial basis of cell identity and castration response in the mouse prostate. Leveraging these data along with a meta-analysis of human prostates and prostate cancer, we identified cellular orthologs and key determinants of ADT response and resistance. Our findings reveal that mouse prostates harbor lobe-specific luminal epithelial cell types distinguished by unique gene regulatory modules and anatomically defined androgen-responsive transcriptional programs, indicative of divergent developmental origins. Androgen-insensitive, stem-like epithelial populations - resembling human club and hillock cells - are notably enriched in the urethra and ventral prostate but are rare in other lobes. Within the ventral prostate, we also uncovered two additional androgen-responsive luminal epithelial cell types, marked by Pbsn or Spink1 expression, which align with human luminal subsets and may define the origin of distinct prostate cancer subtypes. Castration profoundly reshaped luminal epithelial transcriptomes, with castration-resistant luminal epithelial cells activating stress-responsive and stemness programs. These transcriptional signatures are enriched in tumor cells from ADT-treated and castration-resistant prostate cancer patients, underscoring their likely role in driving treatment resistance. Collectively, our comprehensive cellular atlas of the mouse prostate illuminates the importance of lobe-specific contexts for prostate cancer modeling and reveals potential therapeutic targets to counter castration resistance.

**Significance Statement:** Androgen deprivation therapy is a mainstay in prostate cancer treatment, yet many patients eventually develop castration-resistant disease—a lethal progression driven by poorly understood cellular mechanisms. Our study provides a comprehensive cellular map of the prostate, identifying key determinants of normal organization and castration-induced remodeling. By pinpointing the cell types and molecular programs that confer ADT responsiveness or resistance, our findings offer new directions for prostate cancer modeling and pave the way toward novel therapeutic strategies aimed at enhancing ADT efficacy and preventing the emergence of castration-resistant prostate cancer.

## Introduction

Prostate Cancer (PCa) is one of the leading causes of cancer-related death in men worldwide (1). While the disease predominantly manifests in an indolent form, it progresses to an aggressive version in a significant fraction of patients (2). Androgen Deprivation Therapy (ADT) is instrumental in advanced PCa treatment (3, 4). However, inadequate responses and resistance to ADT frequently arises via a plethora of unclear molecular and cellular mechanisms (5) and this state of Castration Resistant PCa (CRPC) is almost always lethal with few therapeutic options. Emerging evidence suggest that epigenetic factors, in addition to or in lieu of genetic mutations, help specific cell types survive through and resist therapeutic insults in PCa (5–8). Unraveling these adaptive cellular responses to ADT is essential for understanding both the emergence and persistence of castration resistance.

Numerous genetically engineered mouse models (GEMMs) have been developed to model both benign and malignant prostatic diseases with varying degrees of penetrance, yet they rarely capture the full progression of human disease (9, 10). Reasons include architectural differences between the mouse prostate and the human counterpart – the former presents as four major (anterior, dorsal, lateral and ventral) lobes (11, 12) and the latter contains three major (central, peripheral and transition) zones (2, 13). Even amidst these major differences, the mouse and human prostate cells are organized within glands and serve the same physiological function (2, 11–13). Specifically, the mouse prostate consists of numerous glands that contain Basal Epithelial cells (BEs), Luminal Epithelial cells (LEs), and Intermediate Epithelial cells (IEs), which are interspersed with rare NeuroEndocrine cells (NEs), and are surrounded by Neuronal cells (Neuro) and FibroMuscular Stroma (FMS) (12). These cells function together to produce components of the seminal fluid, akin to those in the human prostate (12). Therefore, delineating mouse prostate cell type and anatomic equivalents of the human counterpart will enable effective modeling of PCa and other prostatic diseases.

Mouse models of castration effectively recapitulate human responses to androgen deprivation (8, 14–16) and enable precise temporal modeling that is not practical in the human population. Here, LEs are the primary androgen dependent cell types, with castration resulting in widespread LE death and atrophic involution of the prostate (8, 14–16). However, single cell RNA sequencing (scRNAseq) has now shown unexpected diversity in the number of prostate constituent cells (8, 17–22), especially multiple LE subtypes and stem/progenitor cells. Whether these cell types exhibit similar, or distinct androgen sensitivity is under rigorous scrutiny. While the chromatin contexts of these cell types are slowly being understood and have helped with cell phenotyping (18), gene regulatory programs that determine cell identity and castration response remain nebulous. More importantly, whether LE subtypes and stem cells occupy distinct anatomic regions within the multi-lobed prostate and if their location impacts androgen signaling and castration response, is largely unknown. The importance of cellular location is specifically exemplified by prostatic stem cells, whose anatomic location in the proximal or distal prostate is putatively correlated with their distinct roles in development and regenerative outcomes after castration and tissue repair (8, 16–18, 23–25), highlighting potential differences in physiological and pathological contexts. Considering that LEs are posited to be one of the key cell-of-origin of PCa (16, 26), characterizing the transcriptomes, epigenomes and locations of LE subtypes in the normal prostate and dissecting castration responses of these prostatic cells will shed light on drivers of CRPC.

To better understand the cellular constituents of the mouse prostate, their spatial organization and cellular response to castration, we employed an integrated single-cell and spatial omics approach (**Fig. 1A**) that combines scRNAseq, single-cell Multiomics (scMulti), which assesses chromatin accessibility and nuclear RNA from the same cell, and Spatial Transcriptomics (ST). Through this approach we identified cell types, discovered potential gene regulatory modules that drive cell identity, and revealed the organization of these cellular programs in intact prostates (**Fig. 1A**). We find that LE subtypes are enriched in distinct lobes of the mouse prostate and are identified by unique transcription modules (TMs) – i.e. distinct combination of transcription factor and the genes they regulate. Stem cells are enriched in the prostate proximal urethra and the ventral prostate, whilst occurring rarely in other lobes. Although, each lobe-specified LE exhibited distinct androgen responsive transcriptional programs, the ventral prostate contained two types of androgen-sensitive LEs, one bearing semblance to LEs in other lobes and one that is unique to the lobe. Meta-analysis of published scRNAseq data strongly support our findings and integrates multiple datasets to provide a consensus map of the mouse prostate. Comparative analysis of mouse and human prostate scRNAseq data validates cellular concordances and suggests that the ventral prostate appropriately models diverse cell types in the human prostate, informing on future disease modeling. Castration resulted in widespread reorganization of LE transcriptomes and cellular connectivity. These LEs elevated stress responsive (e.g. *Fosl1, Jun, Atf3* and *Nfe2l2*) and stemness associated (e.g. *Klf3/4/5/6* and *Zeb2*) TMs, while dampening Androgen receptor (*Ar*) activity and losing cell-identity specifying TMs (e.g. *Bhlha15*). Strikingly, castration induced TMs from mice were also enriched in tumor cells of ADT-treated and CRPC patients suggesting that cell survival and plasticity programs potentially contribute to the emergence and sustenance of castration resistance. Overall, our cellular cartography effort provides a detailed map of the mouse prostate, highlights the importance of specific lobes in PCa modeling and identifies potential targets to enhance ADT-response and suppress CRPC.

**Fig. 1.**
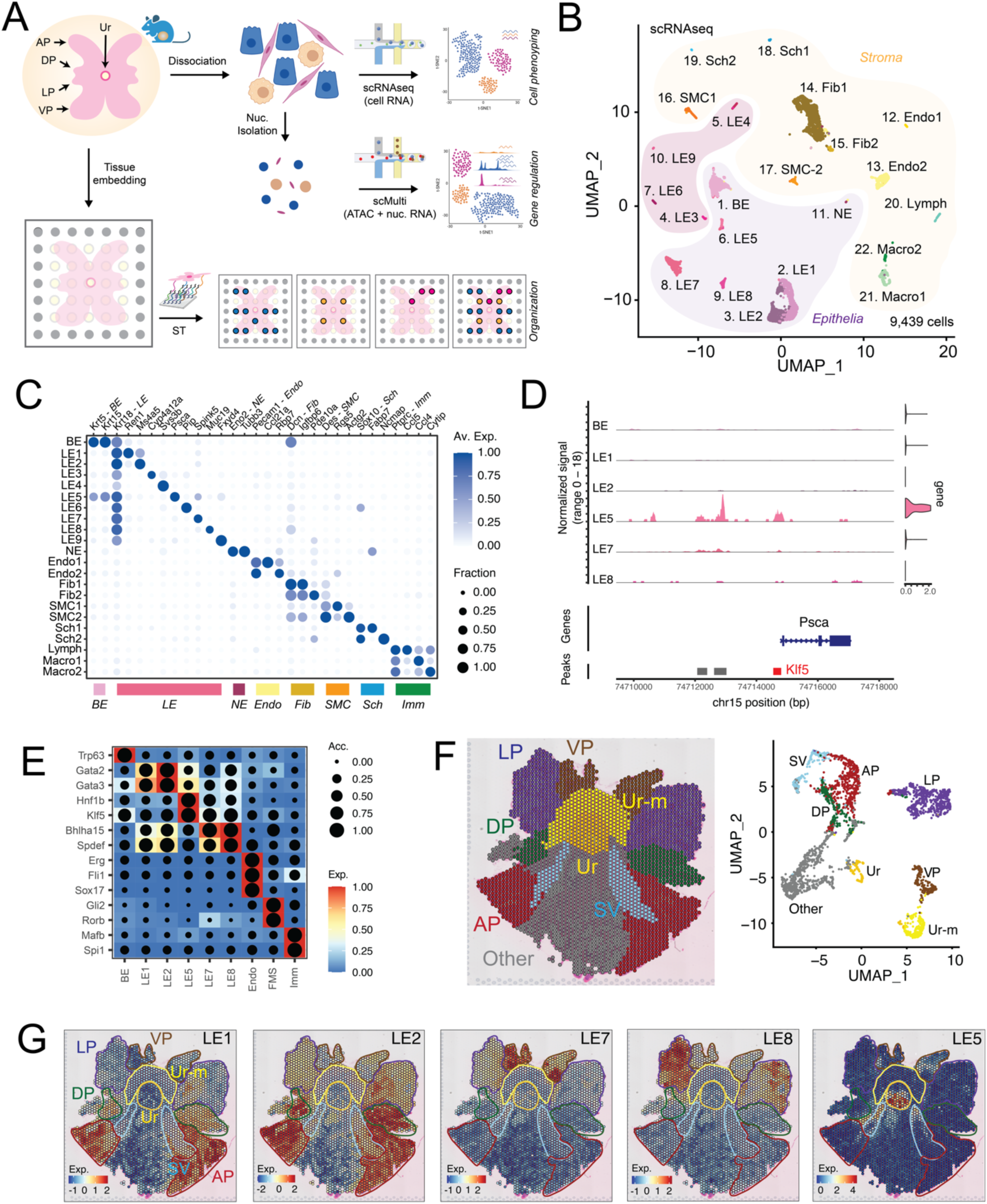
The mouse prostate contains lobe-specific luminal epithelial cells that are represented by unique gene regulatory modules. **(A)** Schematic representing our integrated approach that combines scRNAseq, scMulti and ST analysis to reveal cell phenotypes, gene regulatory features and cellular organization. **(B)** Uniform manifold approximation and projection (UMAP) of individual cells from scRNAseq of 9,439 cells from prostates of C57B6 and FVB mice. Epithelial and NE cell types resembling prior annotations, are highlighted in light purple. Epithelial cell types not resembling prior annotations are highlighted in violet with stromal cells highlighted in yellow. **(C)** Dot plot of gene expression levels in each cell type for selected marker genes with dot size representing fraction of cells expressing the gene and color gradient representing expression levels. **(D)** Coverage plot of Psca across its genomic location, representing ATAC tracks and violin plots of nuclear gene expression in each epithelial cell type. Major aggregate peak locations are represented as grey bars, with red bar highlighting locations enriched for Klf5 motifs. **(E)** Heatmap dot plot showing expression (Exp.) of the eRegulon on a color scale and cell-type specificity of the eRegulon, as indicated by chromatin accessibility (Acc.), on a size scale. **(F)** ST feature plots of histology annotated mouse prostate, with color coded anatomical locations. UMAP of individual spots from ST slide, color coded by anatomical locations. **(G)** ST feature plots with color coded outlines of anatomical locations, representing the expression (Exp.) of LE gene signature (top 20 markers) as a heatmap.

## Results

### Cellular cartography reveals that the mouse prostate contains lobe specific luminal epithelial cells that are defined by unique gene regulatory modules

To characterize cell populations of the mouse prostate, we dissociated individual cells from the entire tissue and subjected them to droplet-based scRNAseq (**Fig. 1A**). We characterized prostates from two commonly used mouse strains in PCa research, namely C57BL/6 (C57B6) and FVB/NJ (FVB). Combined analysis of 9,469 cells from both strains identified 22 distinct cell clusters (see **SI Appendix, Fig. S1A**), with ten *Epcam^+^* epithelial cell types and twelve *Vim^+^* mesenchymal cell types prevalent at almost equal proportions (see **SI Appendix, Fig. S1B-C**). Silhouette analysis supported the uniqueness of each of these cell types (see **SI Appendix, Fig. S1D**). Annotation of cell types based on canonical (prostate) marker genes (**Fig. 1B-C**) showed that our cellular pool contained *Krt5*^+^ BEs, nine types of *Krt18^+^* LEs (LE1-9), *Eno2^+^* rare NEs, two types of *Pecam1^+^* Endothelial cells (Endo; Endo1-2), two types of *Dcn^+^* Fibroblasts (Fib; Fib1-2), two types of *Des^+^*Smooth Muscle Cells (SMCs; SMC1-2), two types of *Sox10^+^* Schwann cells (Sch; Sch1-2) and three types of *Ptprc^+^* Immune cells (Imm) that included Lymphocytes (Lymph) and two types of Macrophages (Macro; Macro1-2). Each of these cell types expressed unique set of marker genes (**Fig. 1C** and see **SI Appendix, Fig. S2A**), underscoring the unique transcriptomes and potential phenotypes of these cells. While a significant majority (seventeen) of these cell types were common across libraires from the two mouse strains, four LEs (LE3, LE4, LE6 and LE9) were predominant in the FVB libraries and one type of Fib was more prevalent in the C57B6 libraries (see **SI Appendix, Fig. S2B-C**). Nevertheless, shared cell types across strains had similar transcriptomes and marker genes (see **SI Appendix, Fig. S1A** and **S2D**).

Prior scRNAseq of prostates from unperturbed and castrated mice had focused on prostatic cells from C57B6 mice (8, 17, 18, 21). To further inform on these studies and to maximize concordance between these published datasets and our work, we chose the C57B6 mice for the rest of the study. scMulti analysis of chromatin accessibility (by Assay for Transposase Accessible Chromatin / ATAC) and RNA from the same nuclei yielded all major cell types identified by scRNAseq, especially diversity in Epithelia (BE, LE1, LE2, LE5, LE7 and LE8) (**Fig. 1A** and see **SI Appendix, Fig. S3A**). For instance, we find that the unique expression of *Psca* (a marker of prostatic stem cells) in LE5 is potentially driven by high accessibility in genomic regions that contain motifs of *Klf5*, a stemness associated Transcription Factor (TF, **Fig. 1D**). Extension of such analysis at the systems level using SCENIC+ (27) revealed that prostatic LEs are defined by unique TMs, wherein the activity of *Gata1/2*, *Klf5*, *Bhlha15* and *Spdef* determine the identity of LE1/LE2, LE5, LE7 and LE8 respectively (**Fig. 1E** see **SI Appendix, Fig. S3B**). Credence to our analysis provided by well validated lineage drivers (such as *Erg*, *Gli2* and *Mafb*) enriched in the appropriate (Endo, FMS and Imm respectively) stromal cell types (28–30) and historically known identifiers of BE, namely *Trp63* (31), arising as a top-hit uniquely in that cell type (**Fig. 1E** see **SI Appendix, Fig. S3B**). In this manner, we were able to attribute cell-specific chromatin accessibility at specific genomic loci to the expression of genes at those loci, rationalizing the expression of BE (e.g. *Sult5a1*), LE1 (e.g. *Mt3*), LE2 (e.g. *Ms4a5*), LE5 (e.g. *Psca and Tacstd2*), LE7 (e.g. *Spink5*) and LE9 (e.g. *Muc19*) transcript markers (**Fig. 1D** see **SI Appendix, Fig. S4A-F**).

We then sought to understand if these diverse cell types are associated with distinct spatial niches in the multi-lobed mouse C57B6 prostate. To this end, we performed ST on whole mount prostate from unperturbed mice (**Fig. 1A, F-G**). Since cells from prostate proximal organs can sometime arise as rare contaminants during surgery and dissociation based single-cell preparations, we used the proximal GenitoUrinary (GU) tract to specifically identify transcriptional and spatial features that are unique to the prostate. Histopathological analysis, based on cellular and glandular morphology (11), enabled the identification of the Anterior Prostate (AP), Dorsal Prostate (DP), Lateral Prostate (LP), Ventral Prostate (VP), Seminal Vesicles (SVs), Urethra (Ur and its muscular surrounding Ur-M) and other nearby GU regions (Other, see **SI Appendix, Fig. S5A**). Strikingly, ST spots from these distinct regions showed up as distinct clusters, indicating large differences in their respective transcriptomes (**Fig 1F**). As expected, *Epcam^+^*spots (indicative of epithelia) and *Vim^+^* spots (indicative of the mesenchyme) and canonical cell type markers (*Krt5, Krt18*, *Eno2*, *Pecam1*, *Dcn*, *Des*, *Sox10*, and *Ptprc*) did not show any regio-specificity within prostate lobes (see **SI Appendix, Fig. S5B-C**). However, mapping scRNAseq derived LE gene signatures showed that LE1 was enriched in the AP, LE2 was enriched in ADP, LE7 was enriched in the VP, LE8 was enriched in the LP and LE5 was enriched in the Ur and VP (**Fig 1G**). Cross-referencing other LEs prevalent in the FVB scRNAseq libraires (LE3, LE4, LE6 and LE9) showed their enrichment in other regions with rare occurrences within prostate lobes (see **SI Appendix, Fig. S5D**). Stomal cell types were spread across the entire tissue section, with SMC2 being mildly enriched in the ADP (see **SI Appendix, Fig. S5E**). Comparative meta-analysis of published scRNAseq datasets support our spatial cell type annotation of LEs and map LE3/4/6/9 to proximal GU regions, namely the SV and the Ejaculatory Duct (ED, **Fig. 2A-F** and see **SI Appendix Table S1**). Put together, we show that the mouse prostate contains lobe-specific LEs that are potentially driven by unique TFs and may correlate with the developmental divergence of primordial cells to drive anatomically different regions of the prostate (12). Our cellular cartography effort, in combination with meta-analysis of public datasets provides a comprehensive reference atlas of the mouse prostate (**Fig. 1-2**, see **SI Appendix, Fig. S1-5** and **Table S1**).

**Fig. 2.**
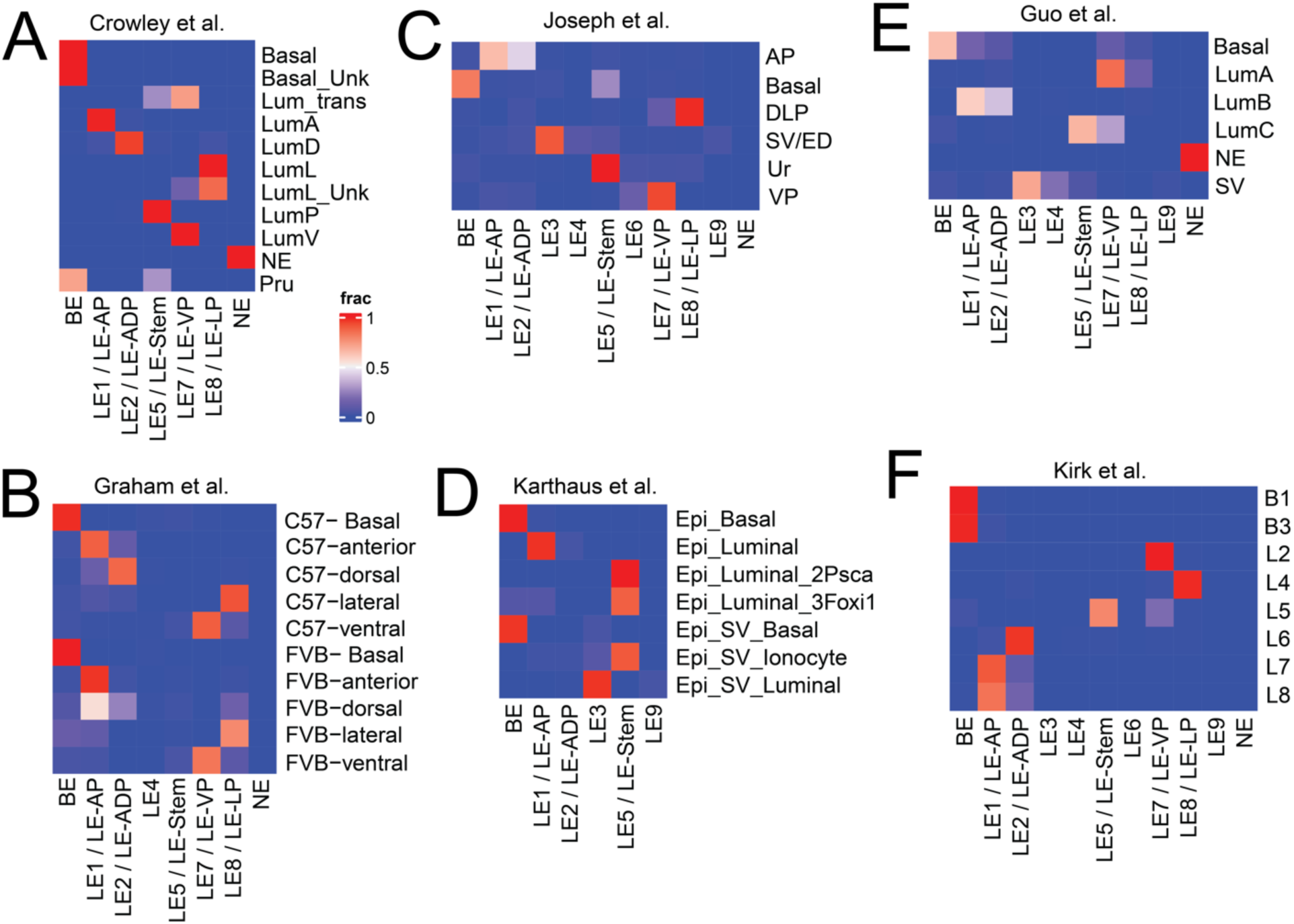
Comparative meta-analysis of mouse prostate single-cell RNA sequencing datasets comparing epithelial cell types identified in this study with published work. (A-F) Heat map representing label transfer analysis of distinct epithelial cell types identified in this study (columns) with cell types annotated in Crowley et al (A), Graham et al (B), Joseph et al (C), Karthaus et al (D), Guo et al (E) and Kirk et al (F).

### The mouse prostate exhibits spatial heterogeneity in androgen responsive gene expression programs

Androgen signaling is crucial for prostate development and homeostasis and is the therapeutic target for androgen dependent PCa (2–4, 12, 14, 15). Therefore, we asked if distinct cell types of the C57B6 mouse prostate expressed a similar set of androgen responsive genes or if each cell type had a unique program. Our scMulti data showed that *Ar*, the core component of androgen responsive gene regulation and driver of PCa, was actively transcribed in all epithelial nuclear populations and in Fib (see **SI Appendix, Fig. S6A**), with steady state cellular RNA levels of *Ar*, as measured with scRNAseq, displaying concordance (**Fig. 3A**). SCENIC+ analysis of scMulti data showed that *Ar* activity, i.e. the extent of chromatin accessibility at Androgen Receptor Elements (AREs) and correlated expression of nuclear *Ar* target gene signature, matched cellular preponderance of *Ar* (see **SI Appendix, Fig. S6A**). In addition, we leveraged canonical Androgen Responsive (And-Resp, Wang et al., see **SI Appendix, Data S1**) gene signatures obtained from bulk omics of mouse prostates (32) and found similar cell-type distributions (see **SI Appendix, Fig. S6A**) and dispersed localization of And-Resp gene set across all prostate lobes (**Fig. 3B**). When we deconvolved the gene signature, we discovered that each prostate lobe had unique sets of androgen responsive genes, with LE1/LE-AP, LE2/LE-ADP, LE7/LE-VP and LE8/LE-LP being marked by *Ren1*, *Plac8*, *Spink1* and *Msmb* respectively (**Fig. 3A-B**). *Pbsn* a standard marker of androgen response and *Ar* activity, was enriched in ADLP, and exhibited significantly lesser expression in parts of the VP (**Fig. 3A-B**). *Gnmt* and *Igfbp2*, LE and SMC enriched And-Resp genes, were dispersed across all lobes (**Fig. 3A-B**). scMulti data showed that *Msmb* and *Plac8* proximal AREs indeed had higher accessibility in LE8/LE-LP, LE1/LE-AP and LE2/LE-ADP and was correlated with increased expression in these specific cell types (see **SI Appendix, Fig. S6B**). Our data supports the notion that distinct lobes activate different sets of androgen response genes.

**Fig. 3.**
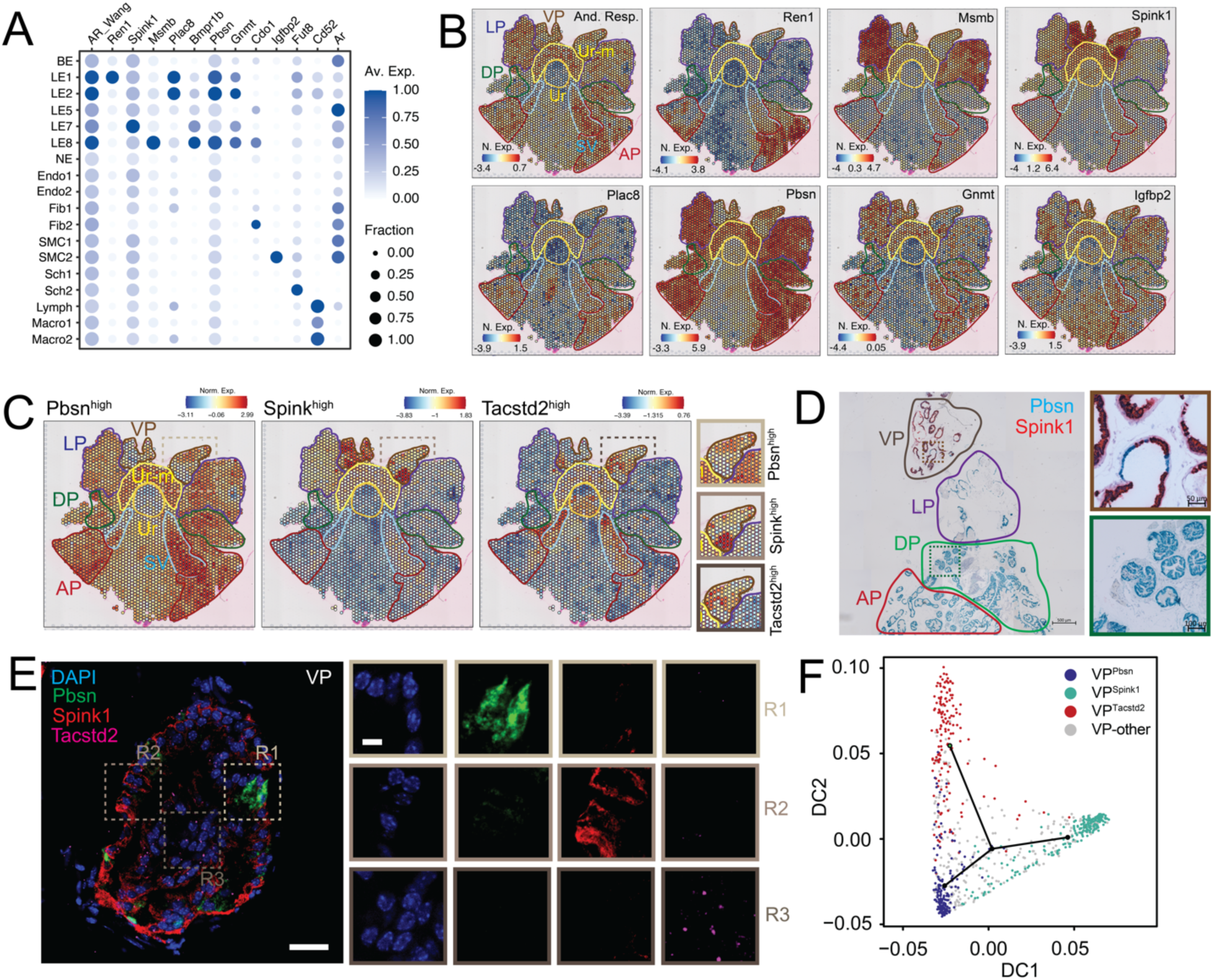
The mouse prostate displays anatomically distinct androgen-responsive and stemness programs, with the ventral lobe enriched for both. **(A)** Dot plot of gene expression levels in each cell type for selected androgen responsive marker genes with dot size representing fraction of cells expressing the gene and color gradient representing expression levels. Combined expression of previously identified androgen responsive (And-Resp) gene set is also shown. **(B)** ST feature plots with color coded outlines of anatomical locations, representing the normalized expression (N. Exp.) of And-Resp gene signature or individual genes as a heatmap. **(C)** ST feature plots with color coded outlines of anatomical locations, representing the normalized expression (Norm. Exp.) of Pbsn^high^, Spink^high^ and Tacstd2^high^ gene signatures as a heatmap. Zoomed-in region of the VP is also shown. **(D)** Representative dual-color RNA-ISH image of a whole mount mouse prostate, with color coded outlines of lobes. Pbsn, blue; Spink1, red. Scale bar, 500 μm. Zoomed-in region of specific glands in the DP (green outline) and VP (brown outline) are also shown. Scale bar, 100 μm, for the DP zoom-in. Scale bar, 50 μm for the VP zoom-in. **(E)** Representative three-color RNA-FISH image of VP localized gland. Pbsn, green; Spink1, red; Tacstd2, magenta; DAPI, blue. Scale bar, 50 μm. Zoomed-in regions of specific areas enriched for Pbsn (R1, light brown outline), Spink1 (R2, brown outline) and Tacstd2 (R3, dark brown outline); are also shown. Scale bar, 10 μm. **(F)** Diffusion plots from trajectory analysis of ST spots representing Pbsn^high^, Spink^high^ and Tacstd2^high^ populations in the VP, i.e. VP^Pbsn^ (blue), VP^Spink1^ (cyan) and VP^Tacstd2^ (red), respectively. Green dot within the plot represents the start of the trajectory. Spots that could not be uniquely annotated as one of these transcriptional programs are in grey.

### The ventral prostate harbors three spatially distinct cell populations: two androgen-responsive cell types and one stem cell population

Upon closer look at the lobular enrichment of distinct And-Resp genes in our ST data, we discovered that a sub-population of VP spots were indeed enriched for *Pbsn* but these spots had minimal VP-enriched *Spink1* expression and *Spink1* expressing spots had low *Pbsn* expression (**Fig. 3B**). Gene sets derived from these *Pbsn^high^* and *Spink1^high^* spots in the VP confirmed that these spatially distinct spots were defined by distinct transcriptional programs (**Fig. 3C**). Strikingly, both these sets of spots had no overlap with spots expressing *Tacstd2/Trop2*, a well validated prostate stem cell marker (33), or *Tacstd2^high^*gene sets (**Fig. 3C**). Expression of these three genes within distinct cells of published VP-specific scRNAseq data (19) supports the existence of distinct *Pbsn^high^*, *Spink1^high^* and *Tacstd2^high^* cell populations (see **SI Appendix, Fig. S7A**). Specifically, *Pbsn* expression within the VP was correlated with *Ren1* (AP enriched), *Msmb* (LP enriched) and *Plac8* (ADP enriched) expression, and anticorrelated with *Spink1* expression (see **SI Appendix, Fig. S7B**), suggesting that *Pbsn^high^* cells in the VP bear semblance to ADLP-specific androgen responsive cells (**Fig. 3C**). Chromogenic RNA *In Situ* Hybridization (RNA-ISH) on whole mount prostates further showed that *Pbsn* and *Spink1* expression are mutually exclusive even within individual glands of the VP, with multicolor and RNA Fluorescence ISH (RNA-FISH) further highlighting *Tacstd2^+^* glandular and peri-glandular cells (**Fig. 3D-E**, see **SI Appendix, Fig. S7C**). SCENIC+ analysis of these distinct sets of spots from ST shows that *Pbsn^high^* cells in the VP (VP^Pbsn^) are identified by *Gata2* TMs, identifiers of LE1/LE-AP and LE2/LE-ADP (**Fig. 1E**), whereas *Spink1^high^* spots (VP^Spink1^) are enriched for *Bhlha15* TMs, potential drivers of LE7/LE-VP (**Fig. 1E**), insinuating additional developmental divergences in the VP (see **SI Appendix, Fig. S7D**). As expected *Tacstd2^high^* spots (VP^Tacstd2^) were identified by stemness associated *Klf5* TMs (see **SI Appendix, Fig. S7D**), identifiers of LE5/LE-Stem (**Fig. 1E**). Trajectory analysis using our ST data (**Fig. 3F**) and published (19) VP scRNAseq data (see **SI Appendix, Fig. S7E**) suggest that VP^Tacstd2^ cells potentially differentiate into VP^Pbsn^ and VP^Spink1^ cells.

### Castration triggers a profound remodeling of epithelial transcriptomes and reshapes cellular interactions

Having mapped cellular locations and androgen signaling across the whole prostate, we sought to understand the impact of castration on these aspects. To address this question, we performed scRNAseq, scMulti and ST of prostates from sham operated (Intact) or orchiectomized (Castrated / Cast) mice (**Fig. 4** and see **SI Appendix, Fig. S8-9**). scRNAseq of 11,976 cells combined from intact and castrated samples showed that the transcriptomes of prostatic epithelia (BE, LE1/LE-AP, LE2/LE-ADP, LE5/LE-Stem, LE7/LE-VP and LE8/LE-LP) were significantly altered by castration (**Fig. 4A**). Expectedly, castration resulted in loss of androgen responsive epithelia and a concomitant gain of stromal, immune cells and androgen insensitive stem cells (LE5/LE-Stem, see **SI Appendix, Fig. S8A**). Analysis of ligand-receptor interactions by CellPhoneDB and CellChat (34, 35) showed dramatic changes in epithelial cell - stromal cell communication after castration, with specific loss of connectivity between the fibromuscular stroma and LE5/LE-Stem or LE7/LE-VP (see **SI Appendix, Fig. S8B**). On the other hand, castration induced increased stromal-epithelial crosstalk is exemplified by *Rspo3-Lgr4*, wherein stromal growth factor (*Rspo3*) secreted communicate with epithelial receptors (*Lgr4*) and this aspect is embedded in distinct spatial patterns (see **SI Appendix, Fig. S8C**). Notably, *Lgr4* expression is relatively higher in the VP in intact conditions but dramatically increases across all lobes upon castration with the highest expression being in the ADP (see **SI Appendix, Fig. S8C**). Diverse types of such transcriptomic reorganization occur - 1) changes in anatomical expression pattern (*Car2* and *Sprr1a,* see **SI Appendix, Fig. S8D**), 2) castration-induced increase in specific lobes (*Cxcl5* and *Slc40a1* see **SI Appendix, Fig. S8E**) and 3) global increase upon castration (*Ly6e* and *Clu*, see **SI Appendix, Fig. S8F**)

**Fig. 4.**
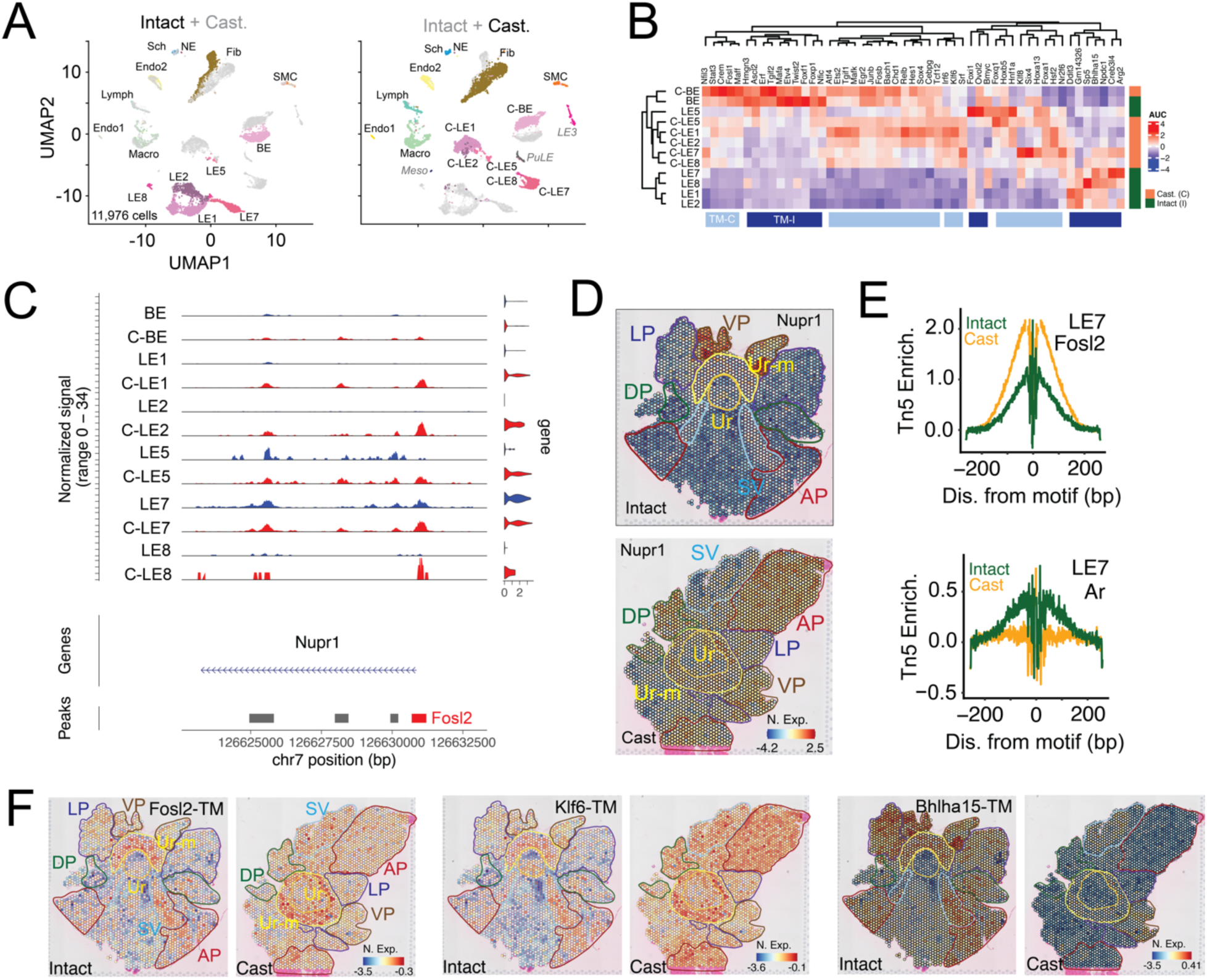
Castration drives dramatic reorganization of cell-specific transcriptomes and induces stress responsive and stemness programs. **(A)** UMAP of 11,976 cells from scRNAseq of prostates from intact (4,958 cells) and castrated (Cast, 7,018 cells) mice. Left, UMAP represents color-coded cell types in the prostate from the intact mice with cell types from the castration samples in grey. Right, UMAP represents color-coded cell types in the prostate from the castrated mice with cell types from the intact mice in grey. Epithelial cell types with dramatic changes in transcriptional programs in the castrated samples are indicated with “C-“ prefix. **(B)** SCENIC-based heatmap of transcription module (TM) enrichment (AUC) in epithelial cells from prostates in intact and castrated mice. Cell types (rows) from prostates of the intact samples are color-coded in green and those from castrated mice are in orange. TMs (columns) enriched in castrated samples are color-coded in light blue (TM-C) and those enriched in the intact samples are color coded in dark blue (TM-I). **(C)** Coverage plot of Nupr1 across its genomic location, representing ATAC tracks and violin plots of nuclear gene expression in each epithelial cell type. Major aggregate peak locations are represented as grey bars. **(D)** ST feature plots with color coded outlines of anatomical locations, representing the normalized expression (N. Exp.) of Nupr1 as a heatmap in whole mount prostates from intact and castrated mice. **(E)** Fosl2 (top) and Ar (bottom) footprints inferred from scMulti ATAC data (Tn5 Enrichment, Tn5 Enrich.) in LE7 cells from prostates of intact (green) and castrated (orange) mice. **(F)** ST feature plots with color coded outlines of anatomical locations, representing the normalized expression (N. exp.) of Fosl2, Klf6 and Bhlha15 TMs as a heatmap in whole mount prostates from intact and castrated mice.

### Prostate epithelial cells respond to castration by inducing stress-response and stemness programs

Dramatic reorganization of transcriptomes upon castration is indicative of cell-intrinsic changes in the epigenome. To dissect specific TMs induced upon castration, we leveraged our scRNAseq and scMulti data (**Fig. 4** and see **SI Appendix, Fig. S9**). SCENIC analysis of our scRNAseq data revealed TMs lost (TM-I) and gained (TM-C) upon castration (**Fig. 4B**). We specifically found that stress responsive TMs such as those driven by *Atf4*, *Fos*, *Fosl1*, *Fosb, Junb*, *Maff*, *Maffk* and *Hsf2* were gained upon castration, along with enrichment of stemness associated TMs, as exemplified by *Klf6*, *Klf8* and *Sox4* (**Fig. 4B**). Motif enrichment within differentially gained ATAC peaks in scMulti data corroborated our findings, highlighting chromatin accessibility gains in AP-1 (*Fos / Jun / Atf*), NFkB, NF1 and KLF family of TFs (see **SI Appendix, Fig. S9A**). As further confirmation of stress-response induction, we find that a well-known stress-induced gene *Nupr1* (36), which is transcribed by AP-1 family of TFs or oxidative stress-activated TF *Nfe2l2*, is expressed in castrated LE cells (C-LEs) with correlated chromatin accessibility gains of *Fosl2* motifs in its promoter (**Fig. 4C**). Consistent with spatial reorganization of transcriptomes, the VP-enriched *Nupr1* is induced in all lobes after castration (**Fig. 4D**). ATAC inferred foot-printing analysis (**Fig. 4E** and see **SI Appendix, Fig. S9B-D**) suggests that stress responsive TFs (e.g. *Fosl2*, *Nfe2l2* and *Atf3*) specifically occupy accessible motifs in LEs and their activity is dramatically increased across all prostate lobes (**Fig. 4F**). *Ar* activity reduced across all epithelia and fibromuscular stroma, as expected, with concomitant loss in lineage/cell-identity driving TMs like *Bhlha15* (LE7/LE-VP, **Fig. 4E-F** and see **SI Appendix, Fig. S9B-C**). Finally, by uniquely leveraging using our scMulti data we generated an expanded Androgen Sensitive (And-Sens, see **SI Appendix, Data S2**) gene set that account for reduction in gene expression in LEs upon castration and concomitant reduction in the accessibility of gene proximal AREs in the same cell (see **SI Appendix, Fig. S9E**). Concordant with loss of *Ar* activity And-Resp and And-Sens gene sets dramatically decreased across all prostate lobes (see **SI Appendix, Fig. S9F**). Notably, And-Resp gene sets are present in proximal GU regions and the prostate, whereas our LE-specific and ARE-correlated And-Sens gene set is restricted to prostatic lobes, enabling the assessment of androgen-mediated *Ar* activity in prostatic epithelia (see **SI Appendix, Fig. S9F**).

### Comparative meta-analysis highlights cell-type parallels between mouse and human prostates and reveals that mouse castration-response programs are enriched in ADT-treated and CRPC patients

Given that the mouse prostate exhibits larger epithelial cell diversity (**Fig. 1-2**) than those currently identified in humans (37), we wanted to identify murine cellular orthologs of the human counterpart. Gene Set Variation Analysis (GSVA) of cellular transcriptomes (**Fig. 5A**) suggests that all LEs in the mouse prostate match with human LE, wherein LE7/LE-VP exhibited the highest semblance. LE5/LE-Stem best recapitulated Hillock and Club cells. When accounting for the cellular heterogeneity in the VP (**Fig. 3**), VP^Pbsn^ and VP^Spink1^ cells matched human LE and VP^Tacstd2^ was similar to Club cells. Meta-analysis of other mouse scRNAseq data corroborates our findings and indicates that the VP accounts for most cell types in the human prostate (**Fig. 5B-D**). We then extended our GSVA to test whether castration in mice effectively recapitulates ADT in humans (**Fig. 5E**). By leveraging published scRNAseq data of ADT-treated and untreated patients (8), we find that human epithelia and tumor cells activate stress responsive and stemness TMs identified in mice. Using published scRNAseq data of CRPC and NEPC patients (6), we find that a significant fraction of these mouse castration response programs are specifically enriched in tumor cells of CRPC patients (**Fig. 5F**). These data suggest that cell survival and plasticity programs significantly contribute to the ADT-response, emergence of castration resistance and its sustenance thereon in PCa patients as well.

**Fig. 5.**
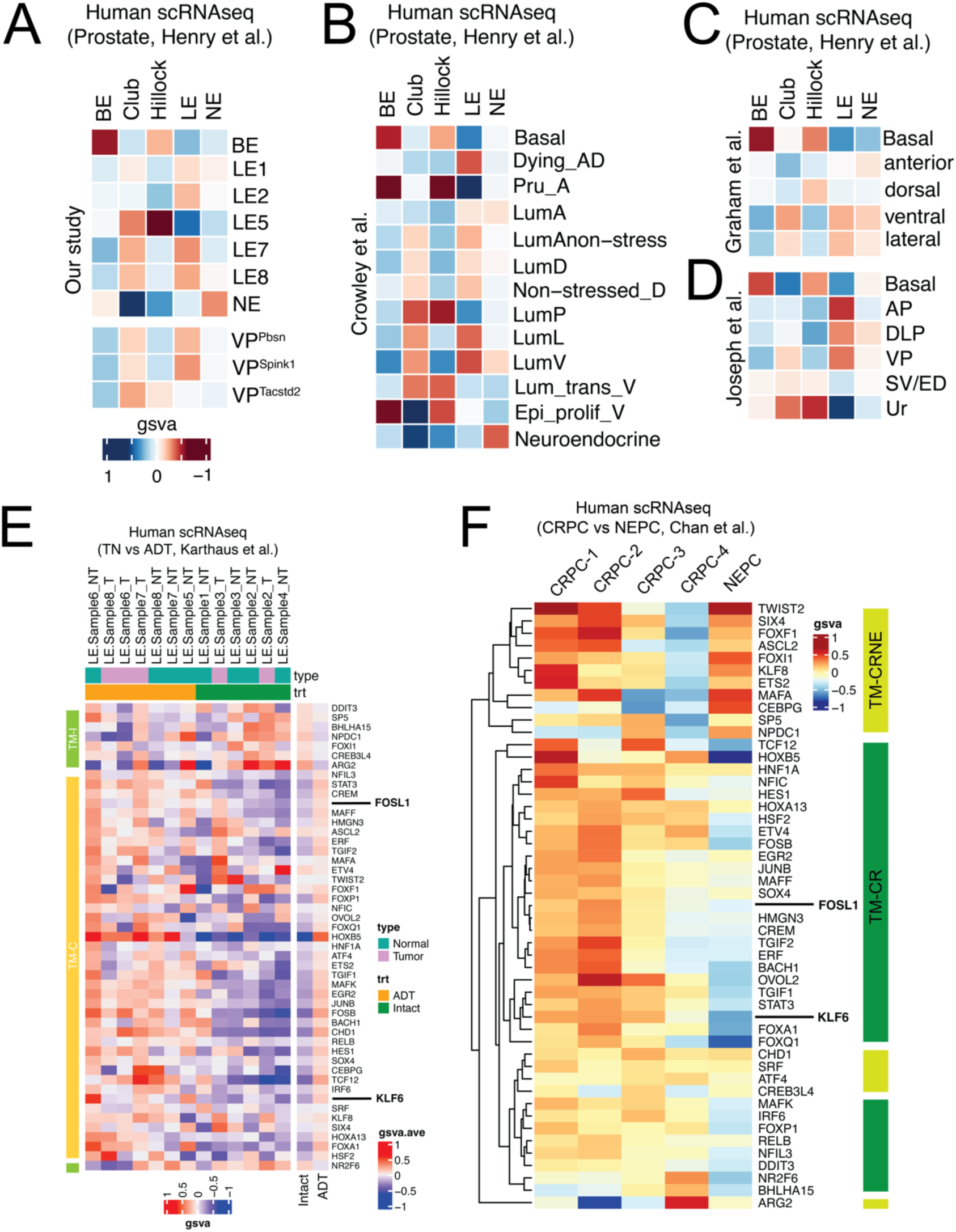
Comparative meta-analysis reveals cell-type parallels between mouse and human prostates and highlights the enrichment of mouse castration-response programs in ADT-treated and CRPC patients. **(A)** Heatmap of gene set variation analysis (gsva) comparing epithelial cell types of the mouse and human prostates. Comparison of distinct cell types identified in the VP (VP^Pbsn^, VP^Spink^ and VP^Tacstd2^) with human prostate epithelia are also depicted. (B-D) Heatmap of gene set variation analysis (gsva) comparing epithelial cell types of the mouse (rows) and human (columns) prostates, wherein mouse prostate data from Crowley et al (B), Graham et al (C) and Joseph et al (D) are represented. **(E)** Gsva based heatmap of TMs in cells annotated as epithelia (cyan, NT) or tumor (magenta, T) from untreated (green, Intact) or ADT-treated (orange, ADT) patients. TMs identified in mouse castration response are represented here. Patient data are from a prior publication. TMs (columns) enriched in cells from untreated patients are color-coded in light green (TM-I) and those enriched in cells from ADT-treated patients are color coded in light orange (TM-C). Averaged gsva for each TM across cell types from untreated (Intact) and ADT-treated patients (ADT) are also depicted **(F)** Gsva based heatmap of TMs in tumor cells from CRPC or NEPC patients. TMs identified in mouse castration response are represented here. Patient data are from a prior publication. TMs (columns) enriched in tumor cells from both CRPC and NEPC patients are color-coded in light green (TM-CRNE) and those enriched in tumor cells from CRPC patients only are color coded in dark green (TM-CR).

## Discussion

ScRNAseq of the mouse prostate has been instrumental in identifying constituent cell types (8, 17–21) and recent scATACseq data has provided chromatin contexts to cellular diversity (18). Our data closely matches these publicly available datasets but additionally provides much needed insights on gene regulatory modules that identify cell types. While correlation of scATACseq datasets with scRNAseq datasets may enable such assessments (18), this approach heavily relies on reference mapping of two distinct zero-inflated single-cell data and thus suffers from low sensitivity. Moreover, this method only correlates chromatin accessibility with steady state RNA in the cell. Access to regulatory modules is uniquely possible only with scMulti, wherein chromatin accessibility is measured along with nuclear RNA levels, indicative of nascent RNA transcription, at individual genomic loci and in the same nucleus. Nevertheless, all these tools require dissociating cells from their native cellular milieu, which can itself induce artifacts (22, 36) and they lack spatial contexts and absolute quantitation of cellular prevalence. We overcame these challenges by further performing ST on whole mount prostates. This aspect is exemplified within our own data pertaining to rare LE3/4/6/9, which seemed to be putative prostatic LEs (*Krt8^+^ / Krt18^+^*) in scRNAseq data. But integrated mapping shows their prevalence in prostate-proximal GU tissues of C57B6 samples, suggestive of these GU cell types being co-enriched during gross surgical dissection of the prostate. However, the rare occurrence of LE3/4/6/9 transcriptomes within intact C57B6 prostate lobes and the presence of these cells only in our FVB prostate library, warrants future spatial and single-cell validations on FVB mice, which have been recently shown to have different transcriptomes than C57B6 (19). Overall, in a first of its kind approach, we integrate scRNAseq, scMulti and ST to perform cellular cartography of the mouse prostate and generate a comprehensive map that informs on cell identity, their gene regulatory determinants their anatomic location.

Our discoveries on epithelial cell diversity are strongly supported by genetic and developmental associations in the prostate. Firstly, we find five LEs (LE1, LE2, LE5, LE7 and LE8) that are identified by distinct gene regulatory modules (*Gata2*, *Gata3*, *Klf5*, *Bhlha15* and *Spdef*) and reside within distinct anatomical locations (AP, ADP, Ur/VP, VP and LP) respectively. Classical IHC has shown Gata2/3 in AP and DLP but absent in the VP (38). Moreover, Gata2/3 knockouts in mice exhibit atrophied prostates with features of BE expansion (38), highlighting their role in LE identity and our data implicates these TFs specifically in LE1/LE-AP and LE2/LE-ADP. Recently, *Klf4* was implicated in maintaining prostate stem cell homeostasis (39), in line with our scMulti data highlighting *Klf* TF family being active in LE5/LE-Stem. *Bhlha15* has been shown to establish secretory cell morphology (40), as expected for LEs, but we have discovered that its activity is enriched specifically in LE7/LE-VP. Developmental associations of prostate derived ETS family member *Spdef* (41) are currently lacking; however our data spurs future interrogations into genetic models that may attribute their importance in LE8/LE-LP. Finally, BEs were enriched for *Trp63* TMs, consistent with the gene being involved in endodermal differentiation to BEs across multiple tissues (42) and its specific necessity in prostate development (31).

Our ST effectively validated lobe-specific scRNAseq data (8, 17–21) and shed light on spatial heterogeneity and quantitative niches within individual lobes of the intact prostate. For instance, previous scRNAseq work had annotated rare occurrences of prostate stem cells in distal ducts of the AP (8, 18, 20), with higher concentration in the proximal prostate and urethra (20, 21). Our integrated analysis supports rarity of these stem cells in the AP (1.5 % of AP-derived ST spots), with similar prevalence found in the DP (3.4% DP-derived ST spots) and LP (2.1% LP-derived ST spots), and high prevalence in the Ur (65.7% of Ur-derived ST spots); however, we discover that these cells are also highly enriched in the VP (33.7% of VP-derived ST spots). Transcriptomes of stem cells across the prostate and the Ur are similar, sans a few markers differentially highlighting ADLVP-(e.g. *B4galnt2*, *Gpr83* and *Cxcl17*) and Ur-stem cells (e.g. *Ly6d*, *Sprr1a* and *Nccrp1*). The location of these cells supports prior reports of both proximal and distal stem cells contributing to prostate regeneration (8, 18). Moreover, the relatively high cellularity of castration insensitive stem cells in the VP also provide a rationale for why this lobe seemed to be least affected by *Ar-* and prostate-disrupting perturbations (38, 43, 44). The VP also contained two distinct androgen sensitive cell types - VP^Pbsn^ and VP^Spink1^, wherein the former cumulatively represented androgen responsive genes that were specific to AP, DP and LP and the latter was unique to the VP. These observations, along with comparative meta-analysis with the normal human prostate suggests that the VP contains diversity of cell types that map to appropriate cell types in the human counterpart (human BE – mouse BE, human LE – mouse LE-VP^Pbsn^ and LE-VP^Spink1^, human Club – mouse LE-VP^Tacstd2^, human Hillock – mouse LE5/LE-Stem and human NE – mouse NE). These observations prompts us to speculate that Pbsn^+^/Spink1^-^ LEs (*Gata2/3*-specific LE1/LE-AP, LE2/LE-ADP and LE7/LE-VP^Pbsn^ or *Spdef*-specific LE8/LE-LP) may be the cell-of-origin for most PRostatic ADenocarcinoma (PRAD), whereas the Spink1^+^/Pbsn^-^ LEs (*Bhlha15*-specific LE-VP^Spink1^) may have a higher predilection to drive SPOP mutant PCa, where human SPINK1 is elevated (45). Moreover, considering that PRAD and Benign Prostatic Hyperplasia (BPH) predominantly arise in distinct zones of the prostate, namely the Peripheral Zone (PZ) and Transition Zone (TZ) respectively (13), our mouse reference map could be used to predict appropriate anatomical matches in the mouse prostate that can model neoplastic and benign diseases.

As expected, castration induced dramatic loss in LE cell-identity (8, 18), denoted by reduction in chromatin activity of lineage specifying TMs such as *Bhlha16* and *Ar*. A recent report suggested loss of LE heterogeneity upon castration (18), whereas we observe retention of heterogeneity. Each LE (LE1/LE-AP, LE2/LE-ADP, LE5/LE-Stem, LE7/LE-VP, LE8/LE-LP) in the intact sample had transcriptionally different counterparts upon castration (C-LE1, C-LE2, C-LE5, C-LE7 and C-LE8) that still expressed a few lobular markers but lost activity of all cell-identity driving TFs. Moreover, we observed reported increases in stromal-epithelial cross-talk (8), as exemplified by increased communication between stromal growth fact *Rspo3* and epithelial receptor *Lgr4*, but did not identify a castration-specific stromal cell population reported in this study (18). It is possible that these differences can be explained by variations in castration time – we focused on early responses (2 weeks) to mitigate effects of cell death, whereas others looked at longer term (4 weeks) responses. Such early castration times may explain why some C-LEs are still lobe enriched, while others are not restricted to any specific lobes - *Car2^+^/Cxcl5^+^*C-LE7 are enriched in the castrated VP, but *Clu^+^/Ly6e^+^* C-LE1 are present in all lobes of the castrated prostate (see **SI Appendix, Table S2**). We additionally find that footprints of *Ar* at accessible AREs do change upon castration. While *Ar* is purported not have a role in BEs (14, 15), our data may provide a reason for *AR* deletion impacting survival and differentiation of BEs (46). Most notably, we observe that stress response transcription programs and stemness programs are induced upon castration in mice and these features are embedded in ADT-treated and CRPC patients. Multiple lines of evidence support the biological and clinical relevance of these programs – 1) CRPC patient derived organoids (named CRPC-WNT and CRPC-SCL) exhibit a combination of our mouse castration response (47), 2) AP-1 family of TFs have been associated with the progression and recurrence of PCa (48), 3) TFs like *Jun* and *Ghrl2* have been reported to drive resistance to hormone therapy in breast cancer (49, 50), while stress-induced TFs such as *Nupr1* have been implicated in the acquisition of chemotherapeutic resistance (51), and 4) Stress responsive TFs, such as *Hsf2*, and stemness factors such as *Tacstd2/Trop2* have been implicated in advanced PCa, with drugs against these targets exhibiting pre-clinical success (52, 53). In summary, our cellular cartography provides a detailed reference map of the mouse prostate, identifies human prostatic orthologs, predicts novel disease modeling approaches, and unveils determinants of castration response and resistance, informing on novel targets for CRPC-therapeutics.

## Methods

### Mouse handling and castration

All animal work was done in compliance with the guidelines of the Institutional Animal Care and Use Committee (IACUC) at University of Michigan. C57BL/6 or FVB mice were purchased from Jackson Laboratory (Bar Harbor, ME). The mice were either sham-operated or castrated at 8 weeks of age. Prostate tissues were harvested from these animals two weeks after surgical procedure.

### Sample collection and tissue processing for scRNAseq

All four prostate lobes were pooled and minced using a pair of spring scissors. Subsequently, the minced tissues were digested with 5mg/mL collagenase Type II (Gibco, 17101015) for 1 hour at 37°C followed by TryLE (Gibco, 17101015) digestion for 10 min. After TryLE digestion, samples were inactivated with an excess of DMEM containing 10% fetal bovine serum (FBS), and samples were sequentially passed through 100 mm and 40 mm cell strainers to remove debris, followed by a 0.5% BSA in 1xPBS wash. Live cells were then pelleted at 300 x g for 5 min at 4 °C. Cell pellets were resuspended in 0.5% BSA in 1xPBS, and counted. Cell suspensions were then processed for scRNAseq by manufacturer’s protocol (10x Genomics, 3’ end assay©) and analyzed using a combination of publicly available and custom packages. Details about library preparation, sequencing and analysis can be found in **SI Appendix, Supplementary methods** and **Data S1.**

### Sample collection and tissue processing for scMulti

Nuclei were extracted from live cell suspensions, as prepared for scRNAseq. Briefly, live cells were pelleted at 300 x g for 5 min at 4 °C. Chilled lysis buffer (10 mM Tris.HCl, 10 mM NaCl, 3 mM MgCl_2_, 1% BSA, 0.1% Tween-20, 0.1% NP-40, 0.01% Digitonin, 1 mM DTT and 1 U/uL RnaseIn, Promega©) was added to cell pellets and incubated on ice for 3-5 min. Chilled Wash Buffer (10 mM Tris.HCl, 10 mM NaCl, 3 mM MgCl2, 1% BSA, 0.1% Tween-20, 1 mM DTT and 1 U/uL RnaseIn, Promega©) was added to the lysed cells, mixed and the nuclei were pelleted at 350 x g for 5 min at 4 °C. Nuclei were washed with Wash Buffer three more times and nuclei pellets were resuspended in Diluted Nuclei Buffer (1x Nuclei Buffer, 10x genomics^TM^, supplemented with 1 mM DTT and 1 U/uL RnaseIn, Promega©) for counting and scMulti library preparation. Nuclear suspensions were then processed for scMulti by manufacturer’s protocol (10x Genomics, ATAC + Gene expression kit ©) and analyzed using a combination of publicly available and custom packages. Details about library preparation, sequencing and analysis can be found **SI Appendix, Supplementary methods** and **Data S2**.

### Sample collection and tissue processing for ST

Mice were orchiectomized and whole prostates were harvested upon removal of adhering fats. Dissected prostates were placed in histocassettes sandwiched between sponges (Fisher©) and fixed in 10% formaldehyde in 1x PBS for 4 h and paraffin embedded within 24 h. FFPE sections of 5 μm thickness were cut by University of Michigan ULAM-IVAC. Whole mount tissues were then processed for ST by manufacturer’s protocol (10x Genomics, Visium Spatial Gene Expression for FFPE©) and analyzed using a combination of publicly available and custom packages. Details about library preparation, sequencing and analysis can be found in **SI Appendix, Supplementary methods** and **Data S3**.

## Data availability

ScRNAseq, scMulti and ST data have been deposited in the Gene Expression Omnibus (GEO), accession number GSE284641, GSE284640 and GSE284571 respectively. All other data are available in the main text or the supplementary information.

## Acknowledgements

We thank Stephanie Miner for reviewing, editing, and submitting this manuscript. We thank Aniket Dagar, Sarah Kang, Nicole Lee and Gregory Raskind for analysis support. The study was supported by Department of Defense Idea Development Award (S.P.), Prostate Cancer Foundation Young Investigator Award (S.P.), NCI Prostate SPORE Career Enhancement Award (S.P.), NCI Outstanding Investigator Award R35CA231996 (A.M.C.), NCI Early Detection Research Network U2C-CA271854 (A.M.C.) and NCI Prostate SPORE grant P50CA186786 (A.M.C.). E.T.K. and spatial transcriptomics were supported by NCI P01 award P010939000. A.M.C is also a Howard Hughes Medical Institute Investigator, A. Alfred Taubman Scholar, and American Cancer Society Professor.

