## Supplementary Information (Methods, Figures and Tables) for "Cellular cartography reveals mouse prostate organization and determinants of castration resistance"

**This PDF file includes:**

Supplementary Methods
Figures S1 to S9
Tables S1 to S2
References

**Other Supporting Information for this manuscript include the following:**

Data S1 to S3 (Excel files)

### Supplementary Methods

***ScRNAseq library preparation and sequencing***

The 10x barcoding and complementary DNA (cDNA) synthesis were performed using 10x chromium 3′ scRNA-seq V2 or V3.1 chemistry according to the manufacturer's instructions. Briefly, an appropriate volume of single-cell suspension/enzyme mix, barcoded beads, and partitioning oil were loaded in distinct lanes on the chip and processed in a 10x chromium controller instrument. The 10x GemCode Technology uses a droplet-based method to partition thousands of cells to generate nanoliter-scale Gel Bead-In-Emulsions (GEMs). Inside each GEM, cDNA generated from an individual cell shares a common 10x barcode. The prepared GEMs were incubated in a thermomixer to produce cDNA. Recovered cDNA was cleaned up and amplified by polymerase chain reaction to generate sufficient quantity for library construction. The size and yield were evaluated using a high-sensitivity bioanalyzer assay chip (Agilent). A set of recommended primers (P5, P7, and R2) in the manufacturer's standard protocol were added to prepare the Illumina-ready sequencing library. Final libraries were evaluated for size and yield using a high-sensitivity bioanalyzer assay chip (Agilent) prior to sequencing with Illumina Hiseq2500 (performed following recommended specifications).

***ScMulti library preparation and sequencing***

Nuclei from mouse prostate tissues were used to generate Chromium Next GEM Single Cell Multiome ATAC + Gene Expression (10x Genomics) libraries following manufacturer's protocol. Briefly, nuclei were prepared from single cells derived from tissue dissociation following the manufacturer's recommended protocol for primary cells as described in CG000365 and nuclei numbers were estimated using Countess II FL automated cell counter. Nuclei transposition were performed for ~6000 targeted nuclei recovery. Following transposition, sequential GEM (Gel Beads-in-emulsion) generation and barcoding steps enables the production of barcoded, full-length cDNA from poly-adenylated mRNA for geneexpression (GEX) library and attachment of a Spacer sequence with barcode to transposed DNA fragments for ATAC library. Post GEM lysis and cleanup, a preamplification PCR step fills gaps and generates sufficient mass for library construction. From this juncture the preamplified product was split to generate ATAC library (where P5 and P7 sequences used for Illumina bridge amplification were added) and GEX library constructions. The GEX library construction step entails enzymatic fragmentation, size selection, and addition of P5, P7, i7 and i5 sample indexes, and TruSeq Read 2 (read 2 primer sequence) via End Repair, A-tailing, Adaptor Ligation, and PCR steps. The generated libraries' size and quality were estimated using BioAnalyzer high sensitivity chip. Multiome ATAC libraries comprise double stranded DNA with standard Illumina® paired-end constructs which begin with P5 and end with P7, Multiome GEX libraries comprise cDNA insert with standard Illumina® paired-end constructs which begin with P5 and end with P7. Sequencing was carried out on an Illumina platform, and data analysis was executed using the 10x Genomics Cell Ranger software, which provided demultiplexing, alignment, and gene quantification against the mm10 mouse genome reference.

***ST library preparation and sequencing***

Serial sections (4x) of whole prostates from sham operated or orchiectomized mice were mounted onto each slide. Tissue sections of each sample type were analyzed to estimate RNA integrity number (RIN) prior to placement on Spatial Tissue Optimization slides. Optimization was performed only for samples with RIN greater than or equal to 7. Spatial gene expression slides specific for either cryosections or FFPE sections were used (10X Genomics™) and were submitted to the Advanced Genomics Core (AGC, University of Michigan) for tissue optimization, permeabilization and library construction. Tissue was processed and placed according to the manufacturer's instructions (CG000408 Visium Spatial Gene Expression for FFPE –Tissue Preparation Guide and CG000407 Visium Spatial Gene Expression for FFPE User Guide). Samples were stained lightly with H&E on the Spatial Gene Expression slides using a standard protocol (CG000409 Visium Spatial Gene Expression for FFPE – Deparaffinization, H&E Staining, Imaging & Decrosslinking). H&E images were scanned at high resolution and then probed with the Visium Mouse Transcriptome Probe Set (Visium Mouse Transcriptome Probe Set v1.0). Sequencing depth for probe set assay was 100 million reads/sample. All sequencing was performed at the University of Michigan AGC on the NovaSeq6000 with 300 cycles and 150 bp paired end reads. Each sequenced library was demultiplexed to FASTQ files and aligned to reads to the mm10 mouse transcriptome with the Spatial Space Ranger 1.3©.

***In situ hybridization***

For the in-situ validation RNA in situ hybridization (RNA-ISH) was performed as single-plex, duplex chromogenic assay and for cross validation they were performed as three-plex RNA fluorescence in situ hybridization (RNA-FISH) assay. The RNAscope 2.5 HD Brown assay from Advanced Cell Diagnostics was used to perform RNA-ISH targeting the Mm-Spink1 gene (catalog no. 532311) and Mm-Probasin (catalog no. 30031). For dual RNA-ISH, the RNAscope 2.5 HD duplex assay was employed with probes for Mm-spink1, and Mm-probasin (catalog no. 409091, and 532311-C2). RNA quality for single-plex was checked using a positive control probe for the housekeeping gene Mm-Ppib (catalog no. 313911) and for duplex Mm-Ppib in C1 and Mm-Polar2 in C2 (catalog no. 321651), while assay background was assessed with a negative control probe for the bacterial DapB gene (catalog no. 322000). Each RNA molecule appeared as a distinct brown dot under the microscope as single-plex and as red/green for Mm-spink1 and Mm-Probasin respectively for the duplex assay. For simultaneous detection of three genes (three-plex) we employed RNAScope Multiplex fluorescent reagent kit V2 employing the probes for Mm-Probasin, Mm-Spink3 in channel2 (C2), and Mm-Tacstd2 in channel 3 (C3)- (catalog no. 526301, 409091-C2, and 471751-C3). Like the chromogenic assays, positive and negative assay control probes were used. For positive probe 3-plex Positive Control Probe: Mm-RNAscope mouse positive control probe for RNAscope Multiplex Fluorescent Assay -Polr2a (C1 channel) and PPIB (C2 channel), UBC (C3 channel) was used and for negative probe: RNAscope Negative control probe DapB (of Bacillus subtilis strain) for RNAscope Multiplex Fluorescent Assay was used. To develop the corresponding colors TSA Vivid dyes were used employing fluorophores 520, 570, and 650 (catalog no. 323271, 323272, and 323273). Three study pathologists independently evaluated the staining at ×100, ×200, and ×400 magnifications to determine the presence and patterns of RNA expression.

***ScRNAseq data preprocessing***

After sequencing, read demultiplexing, alignment, and gene quantification were performed using the 10X Genomics Cell Ranger pipeline (v5.0) with the pre-built reference genome (mm10 for mouse). Unless otherwise specified, downstream analyses were conducted using the R package **Seurat** (v4.1) (1) on the filtered gene count matrix. Low-quality cells were further filtered based on total UMI counts (nCount_RNA), the number of detected genes (nFeature_RNA), and the percentage of mitochondrial reads (percent.mt) per cell. Outliers were identified using the isOutlier function from the **scater** package (2), with thresholds set at 3 median absolute deviations (MADs) from the median for both total UMI counts and detected genes, and higher outliers or percentages $\geq$ 80% for mitochondrial reads. Specifically, cells flagged as outliers in any of these metrics were removed (3). Following cell filtering, mitochondrial genes were excluded from the matrix. The resulting count matrix was normalized using the NormalizeData function with the "LogNormalize" method. Highly variable genes (HVGs) were identified using the FindVariableFeatures function with the “vst” method, followed by dimensionality reduction steps including ScaleData, RunPCA, and RunUMAP, to obtain a two-dimensional map of the cells. Cells were assigned to clusters using the FindNeighbors and FindClusters functions. Cluster markers were identified using FindAllMarkers and visualized with DotPlot. For data pooled from multiple libraries, batch correction was applied using the **scran** (4) and **batchelor** (5) packages. HVGs across libraries were identified using modelGeneVar with the "block" option, and the getTopHVGs function was used to select HVGs. Batch effects were corrected using fastMNN based on the selected HVGs before generating the UMAP.

***ScRNAseq data analysis and annotation***

Cell populations in the dataset were initially annotated based on the expression of epithelial (EPCAM) and mesenchymal (VIM) cell markers, resulting in the identification of 10 epithelial and 12 mesenchymal clusters. Further characterization of these clusters was performed using canonical markers, such as Keratin 5 (Krt5) for basal cells and Keratin 18 (Krt18) for luminal cells. This enabled the classification of nine distinct luminal epithelial cell types, one basal type, a neuroendocrine type, and 11 stromal cell types. Validation of the clustering results was conducted using the **Buster** package (v1.13.0), which calculated silhouette scores to assess cluster robustness. Unique markers for each cell type were identified using the **Seurat** package (v4.1), and the results were visually represented in a bubble plot. To align cell type annotations with previously published datasets (6-9), a label transfer approach was applied. Cell type annotations from our dataset were used as a reference, and annotations were transferred to the published datasets. This process involved identifying anchor points between the reference and query datasets using the FindTransferAnchors function, followed by TransferData to assign predicted cell types to the query data. To ensure high confidence in the transfer, only annotations with a prediction score above a defined threshold (prediction.score.max > 0.8) were considered. The resulting proportion of cells in each predicted cell type was calculated and visualized in a heatmap to identify the best-matched cell types between the reference and query datasets. This process generated a consensus table summarizing the alignment of cell types across datasets, providing a clear basis for further comparative analyses.

***Transcriptional module data analysis with SCENIC***

Transcriptional module analysis was conducted using **SCENIC** (v1.2.2) with R (v3.6.3), following a detailed tutorial provided by SCENIC (https://rdrr.io/github/aertslab/SCENIC/f/vignettes/SCENIC_Running.Rmd). The process began with the initialization of a settings configuration using the initializeScenic function. Next, the runGenie3 function was employed to generate the co-expression network, forming the foundation for the computation of the Gene Regulatory Network (GRN). Subsequent steps involved the use of SCENIC functions: runSCENIC_1_coexNetwork2modules to define co-expression modules, runSCENIC_2_createRegulons to identify regulons, and runSCENIC_3_scoreCells to score cells based on regulon activity. Promoter information from the "10kb" dataset was referenced during this process. The computed GRN data were retrieved using the loadInt function, and the CoexWeight value was utilized as the derived metric from the SCENIC computation.

***ST data processing and analysis***

Raw sequencing data was processed using the 10x Genomics Space Ranger pipeline(v1.3) to generate FastQ files. The sequences were aligned to the mouse mm10 genome, and gene expression counts were quantified using the default settings of the Space Ranger pipeline. Expression levels of RefSeq coding genes were also quantified using the pipeline's default parameters. Mouse prostate sections were prepared and analyzed using the **10X Visium** platform. The regions of interest within the 10X Visium sample were manually annotated to delineate the lobes. These manual annotations were subsequently converted into image annotations using the **Loupe Browser**. Unless otherwise specified, downstream analyses were conducted using the **R package Seurat** (v4.1). The average log-transformed expression of the top 20 marker genes for each cell type, derived from both the current and previously published single-cell RNA-seq datasets, was then mapped onto the spatial image for visualization. Additionally, expression data were normalized using ACTB as a reference to facilitate the comparative visualization of gene expression changes between intact and castrated conditions. This normalization helped highlight physiological alterations in response to castration across different anatomical regions.

***Subgrouping VP in ST and public scRNAseq dataset***

To investigate the ventral prostate subgroups and their trajectories ST and scRNA-seq data (7) were utilized. Each dataset was preprocessed by normalizing gene expression data, identifying highly variable genes, and performing principal component analysis (PCA). For spatial transcriptomics datasets, the **SCT assay** was used, while the **RNA assay** was employed for scRNA-seq datasets. Distinct cell subgroups, including "spink1," "pbsn," and "tacstd2," were identified based on normalized gene expression levels (log-transformed). To define these subgroups, the top 25% of cells with the highest expression levels for each marker gene were retained, ensuring high specificity. Subgroups without significant marker expression were labeled as "ventral_rest." These subgroups were further refined using a combination of cluster-level annotations. To visualize the VP subgroups in UMAP, integration across samples within datasets was achieved using Seurat's anchor-based integration workflow. Features were harmonized across samples using **SelectIntegrationFeatures** and

**FindIntegrationAnchors**, followed by data integration using **IntegrateData** with adjusted weight parameters to account for sample variability. Dimensionality reduction was performed using **UMAP**.

***Trajectory analysis***

Trajectory analysis was conducted using diffusion maps and slingshot-based lineage reconstruction. Dimensionality reduction was performed using **DiffusionMap** from the **destiny** (v3.8.1) package, followed by trajectory inference with **slingshot** (v2.2.1). Key lineages were identified by setting the starting cluster as "Tacstd2" to trace trajectories across VP subgroups. The reconstructed trajectories and lineage structures were visualized with scatterplots, color-coded by cell subgroups for both spatial transcriptomics and scRNA-seq datasets.

***Mouse-Human cell type mapping***

To investigate the correspondence between mouse and human prostate cell types, scRNAseq data from in-house datasets, published datasets (6, 7, 10), and ST data were utilized. Mouse gene symbols were converted to their human homologs, and only genes with valid HGNC symbols were retained. The converted dataset was processed into a **Seurat** object, normalized, and filtered for highly variable genes. The analysis focused on epithelial subgroups, including Basal, Luminal, and Neuroendocrine (NE) cells, with the top 50 marker genes for each subgroup identified using **Seurat**'s **FindAllMarkers** function. Gene set variation analysis (**GSVA**) was conducted to assess the enrichment of mouse epithelial marker genes in human prostate datasets, using log-transformed TPM expression data. The **GSVA** results were normalized and visualized in a heatmap, where rows represented mouse cell types and columns represented human cell types, highlighting conserved transcriptional similarities across species.

***Ligand-receptor analysis***

We performed a differential ligand-receptor analysis on the annotated single-cell data using CellphoneDB (11). The data were first separated between intact and castrated, and re-normalized using the NormalizeData function with the "LogNormalize" method. In each condition, we calculated the differentially expressed genes for each cell annotation using FindAllMarkers and selected those with average log2 fold-change greater than 0.1 and an adjusted p-value of less than 0.05. The normalized data, cell annotations, and differential genes were used as inputs to CellphoneDB's DEG_analysis function which returns all relevant interactions. These are defined as interactions where all the genes in the interaction are expressed in more than 10% of the cells of interest and one of the genes in the interaction is included in the differentially expressed genes. Differential ligand-receptor interactions were defined as gained, if the interaction was not relevant in the intact subset but was relevant in the castrated subset, and lost, if the interaction was relevant in the intact subset but was not relevant in the castrated subset. We further validated our analysis using another cell-cell interaction toolkit, CellChat (12) and their netVisual_diffInteraction method.

***Multiome data analysis***

Following sequencing, raw reads were processed with the 10X Genomics Cell Ranger ARC pipeline (version 2.0). Filtered RNA count matrix and ATAC fragments were used as input for downstream analyses with Signac (version 1.5.0) (13). SoupX (4) was used to correct the RNA count matrix. After visualizing QC metrics (total UMIs per nucleus, total fragments per cell and TSS enrichment) with violin plots, outlier nuclei were removed. In addition, nuclei with more than 40% mitochondrial reads were removed. We noticed that although more nuclei were called for the castrated prostate library, relatively lower quality in RNA (median UMIs per cell and median genes per cell) was seen. Therefore, an equal number of nuclei as the intact library were kept to match RNA quality of both libraries. After QC, peaks calling was redone with MACS2 (14) using the callPeaks function. Cell annotation was based on label transfer using annotated scRNA-seq libraries from intact mouse prostate as reference. Dimension reduction (UMAP) of the RNA data was based on top 2000 HVGs and the first 30 principal components; dimension reduction of the ATAC data was based on the top 25% peaks and the top 30 latent semantic indexing (LSI) components except the first component because it was highly correlated with sequencing depth. To find differentially accessible regions between groups of nuclei, the findMarker function was utilized with the LR (logistic regression) test method and the total number of fragments as a latent variable. Cell level motif activity score was computed by running chromVAR (15) with HOMER motifs (16). Motif enrichment scores for peaks around genes of interest were calculated using motifmatchr (17). Heatmap of ATAC signals in motif binding sites was generated with deeptools (18); motif binding sites were first identified with HOMER script scanMotifGenomeWide.pl to obtained enrichment per cell type, bam files of ATAC read alignment for each cell type were extracted from the whole library bam file using corresponding list of nuclear barcodes. Footprinting analysis (13) was used to estimate the extent of transcription factor binding based on ATAC signals (19). Regulon analysis by integrating gene expression and ATAC of the intact multiome samples was performed with SCENIC+ (v0.1) (20).

### Supplementary Figures

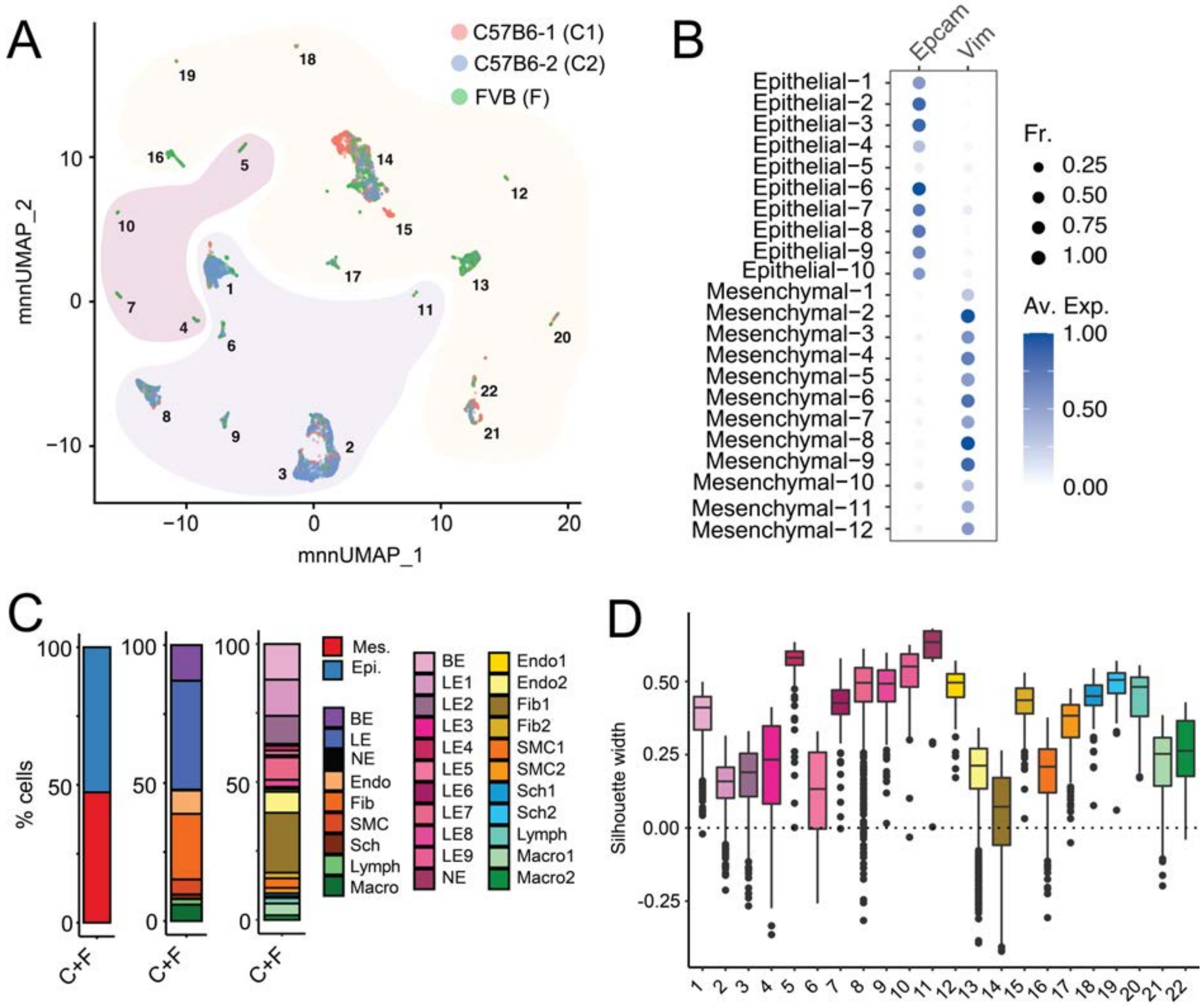


**Fig. S1. Related to figure 1. Benchmarking mouse prostate scRNAseq data. (A)** Batch corrected (mnn) UMAP of 22 distinct cell types from scRNAseq of 9,439 cells from prostates of C57B6 and FVB mice, with cells being color coded by sample. C57B6-1 (C1) and C57B6-2 (C2) are two different sample sets of 4 prostates each, procured, processed and sequenced on distinct days. **(B)** Dot plot of Epcam and Vim gene expression levels in each cell type with dot size representing fraction of cells expressing the gene and color gradient representing expression levels. **(C)** Stacked bar plot of the percentage of cell types, color coded by epithelial vs mesenchymal cells, histologically known mouse prostate cell types and cell types discovered in this study respectively. **(D)** Bootstrapped silhouette analysis depicting silhouette widths for each cell type.

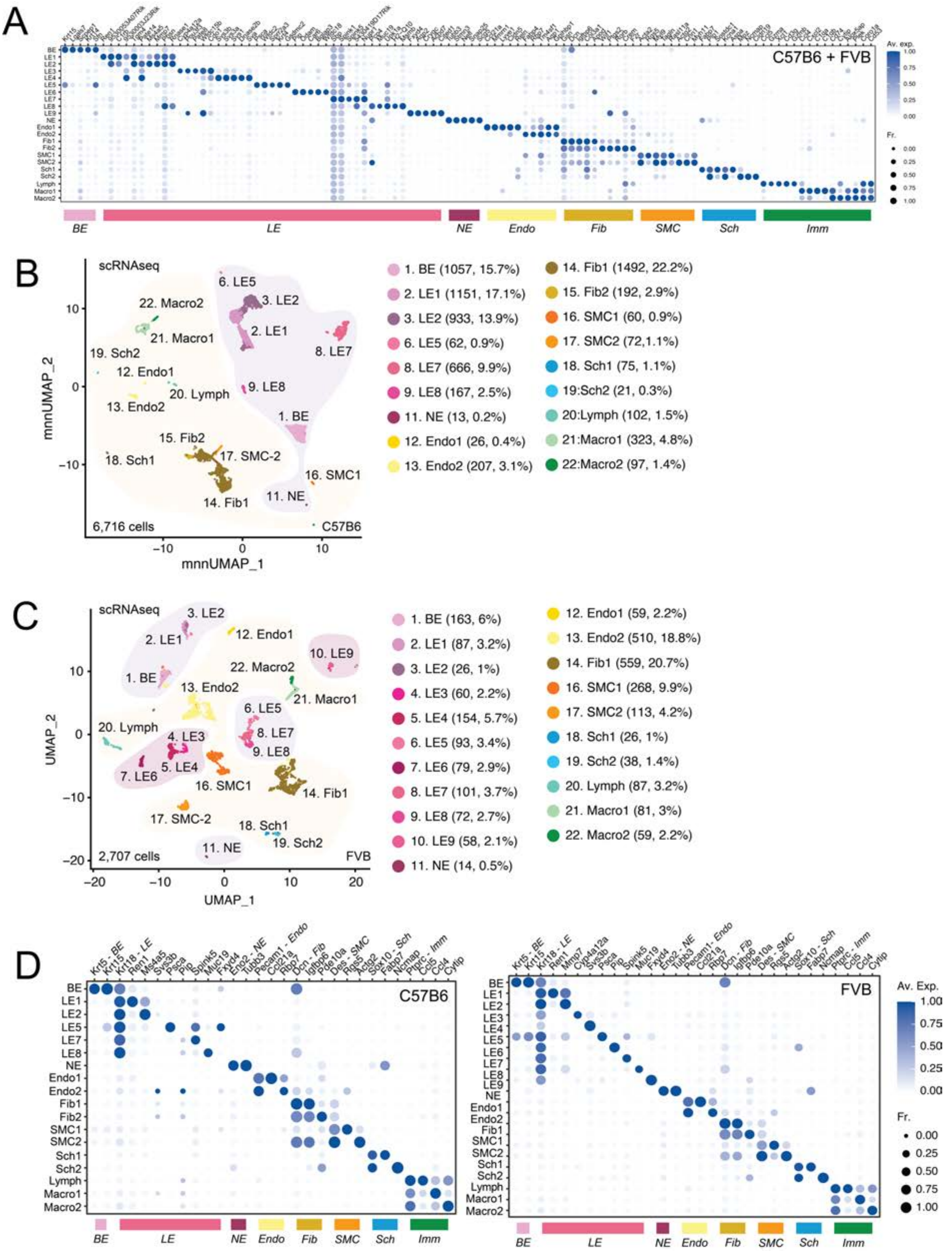


**Fig. S2. Related to figure 1. Cell type identification in distinct mouse strains. (A)** Dot plot of gene expression levels in each cell type (rows) for selected (top five) marker genes (columns) with dot size representing fraction of cells expressing the gene and color gradient representing expression levels in prostates from C57B6 and FVB mice combined. **(B)** Batch corrected (mnn) UMAP of 18 distinct cell types from scRNAseq of 6,716 cells from prostates of C57B6 mice. Cell type color codes and numbers are set to match those in Fig. 1B. Number of cells and percentage prevalence of cells are also depicted.

**(C)** UMAP of 21 distinct cell types from scRNAseq of 2,707 cells from prostates of FVB mice. Cell type numbers are set to match those in Fig. 1B. Cell type color codes and numbers are set to match those in Fig. 1B. Number of cells and percentage prevalence of cells are also depicted. **(D)** Dot plot of gene expression levels in each cell type (rows) for selected marker genes (columns) with dot size representing fraction of cells expressing the gene and color gradient representing expression levels in prostates from C57B6 mice (left) and FVB mice (right).

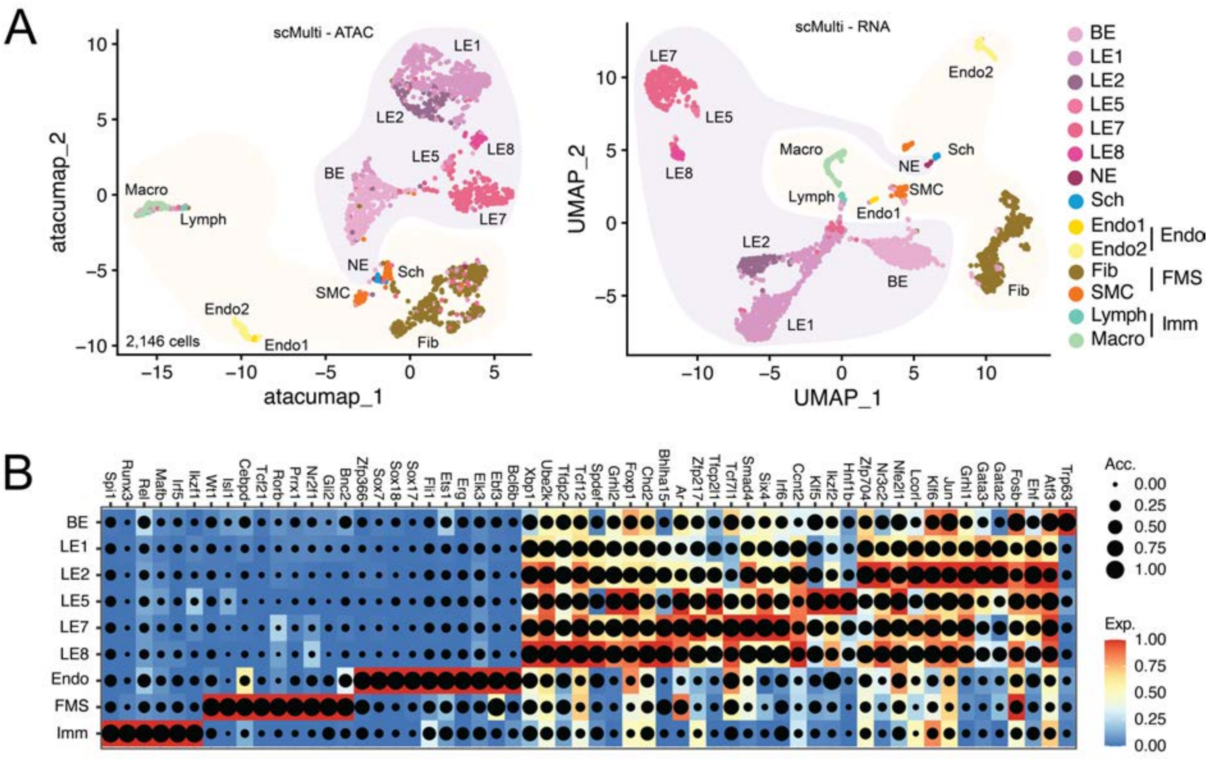


**Fig. S3. Related to figure 1. Cell type mapping in scMulti data. (A)** UMAP of 14 distinct cell types from scMulti of 2,146 cells from prostates of C57B6 mice. Cell type color codes are set to match those in Fig. 1B. Epithelial and NE cell types resembling prior annotations, are highlighted in light purple and stromal cells highlighted in yellow. Left, UMAP based on ATAC data and right, UMAP based on RNA from the same nuclei. **(B)** Heatmap dot plot showing expression (Exp.) of the eRegulon on a color scale and cell-type specificity of the eRegulon, as indicated by chromatin accessibility (Acc.), on a size scale. Endo1 and Endo 2 cells from (A) were pooled to Endo, Fib and SMC from (A) were pooled to FMS and Lymph and Macro from (A) were pooled to Imm to robustly compare TMs of stroma and epithelial cells.

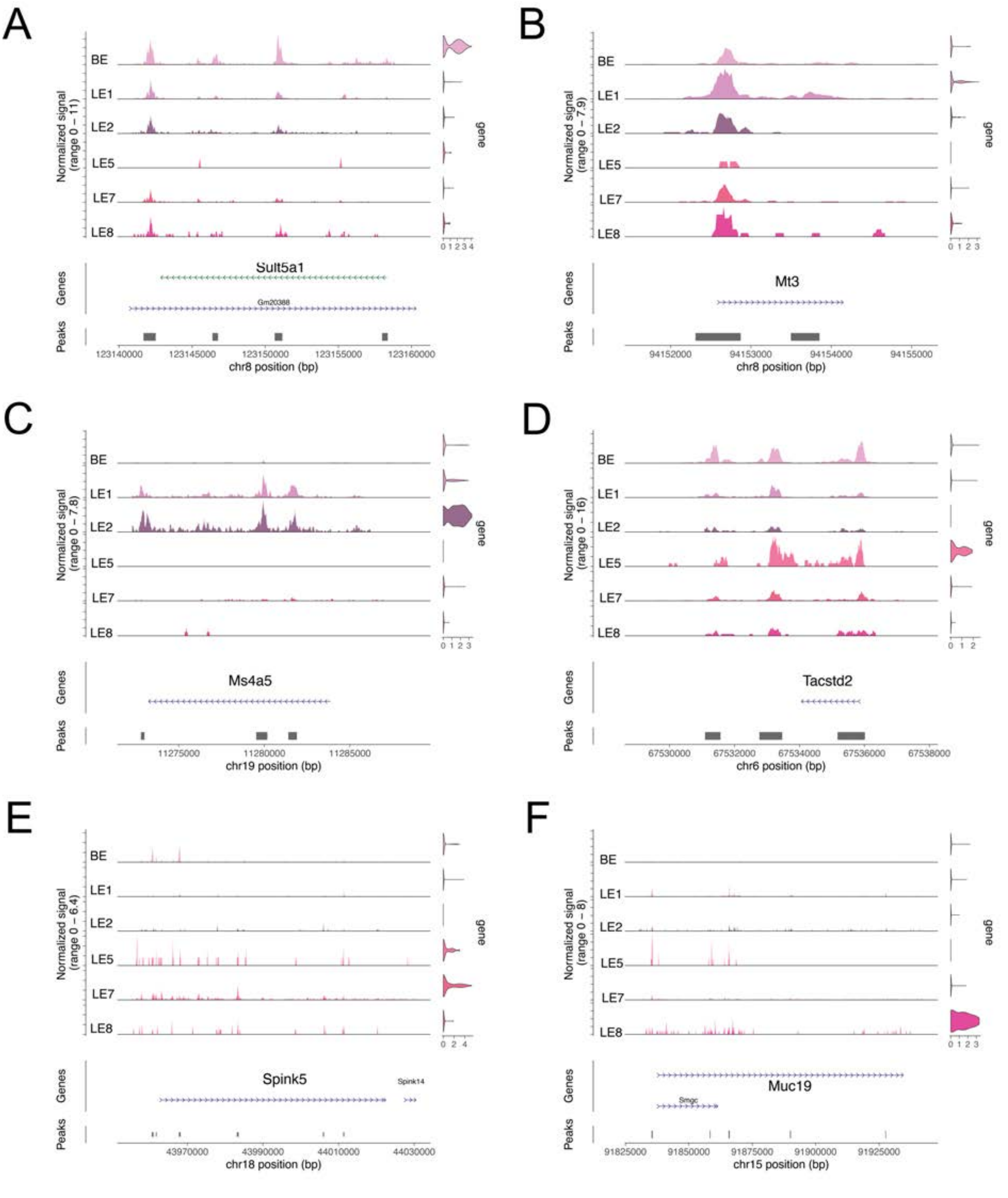


**Fig. S4. Related to figure 1. Cell type specific gene expression and their appropriate ATAC peaks. (A-F)** Coverage plots of *Sult5a1* (BE-specific, A), *Mt3* (LE1 specific, B), *Ms4a4* (LE2 specific, C), *Tacstd2* (LE5 specific, D), *Spink5* (LE7 specific, E) and *Muc19* (LE8 specific, F) across their respective genomic locations. ATAC tracks and violin plots of nuclear gene expression in each epithelial cell type is represented. Major aggregate ATAC peak locations are represented as grey bars.

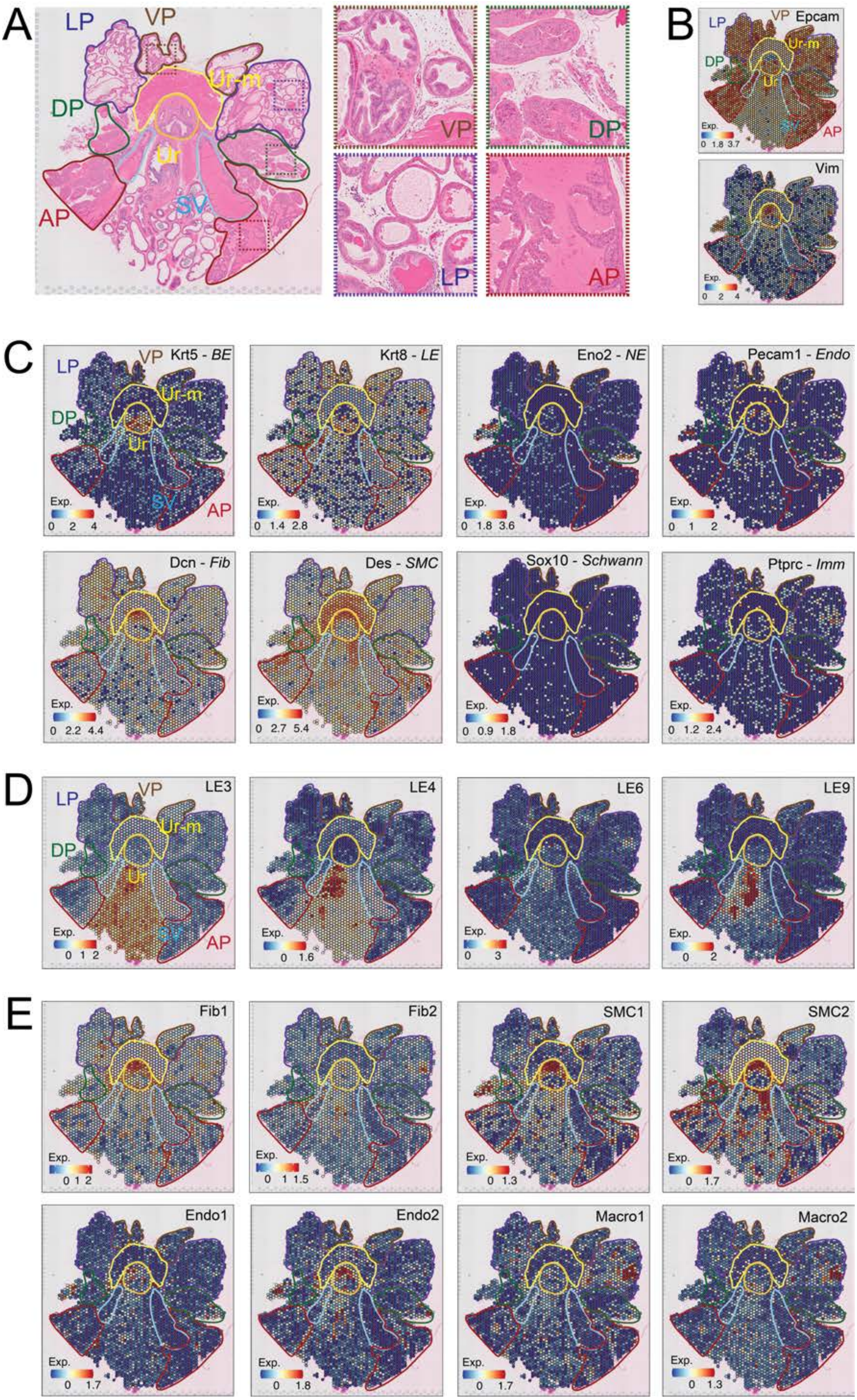
A
LP
VP
Ur-m
DP
Ur
AP
SV
B
Epcam
Vim
Exp.
C
Krt5 - BE
Krt8 - LE
Eno2 - NE
Pecam1 - Endo
Dcn - Fib
Des - SMC
Sox10 - Schwann
Ptprc - Imm
D
LE3
LE4
LE6
LE9
E
Fib1
Fib2
SMC1
SMC2
Endo1
Endo2
Macro1
Macro2

**Fig. S5. Related to figure 1. Mapping of specific genes and cell-specific gene expression programs to ST data. (A)** H&E image of whole mount mouse prostate on the ST slide with color coded anatomical locations annotated based on histological analysis. Zoomed in regions of representative glands of AP, DP, LP and VP are shown. **(B)** ST feature plots with color coded outlines of anatomical locations, representing the expression (Exp.) of *Epcam* and *Vim* genes as a heatmap. **(C)** ST feature plots with color coded outlines of anatomical locations, representing the expression (Exp.) of canonical prostate cell type markers, namely, *Krt5* (BE), *Krt8* (LE), *Eno2* (NE), *Pecam1* (Endo), *Dcn* (Fib), *Des* (SMC), *Sox10* (Schwann / Neuro) and *Ptprc* (Imm), as a heatmap. **(D)** ST feature plots with color coded outlines of anatomical locations, representing the expression (Exp.) of epithelial cell signatures not resembling prior annotations, i.e. LE3, LE4, LE6 and LE9 gene signatures, as a heatmap. **(E)** ST feature plots with color coded outlines of anatomical locations, representing the expression (Exp.) of stromal cell signatures as a heatmap.

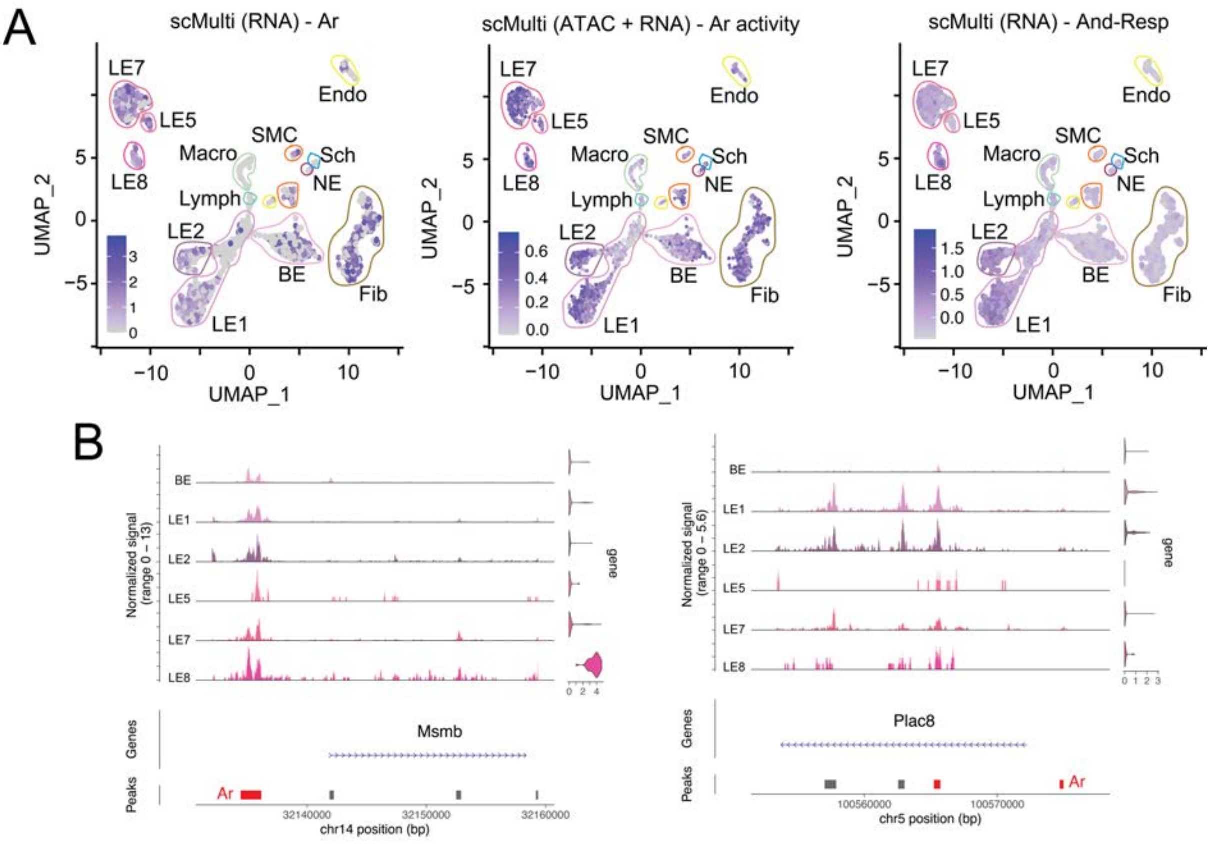


**Fig. S6. Related to figure 2. Ar is expressed and active in epithelial and stromal cell types, but distinct cell types express different target genes. (A)** UMAP of 14 distinct cell types from scMulti of 2,146 cells from prostates of C57B6 mice, with color coded cell cluster outlines based on Fig 1B. Left, Expression of *Ar* is represented as a feature on this UMAP, with the expression depicted as a heatmap. Middle, SCENIC-based *Ar* activity is represented as a feature on this UMAP, with the activity score depicted as a heatmap. Right, Expression of androgen responsive gene set (And-Resp, Wang et al) is represented as a feature on this UMAP, with the expression depicted as a heatmap. **(D)** Coverage plots of *Msmb* (left) and *Plac8* (right) across their genomic locations, representing ATAC tracks and violin plots of nuclear gene expression in each epithelial cell type. Major aggregate peak locations are represented as grey bars, with red bars highlighting locations enriched for AREs.

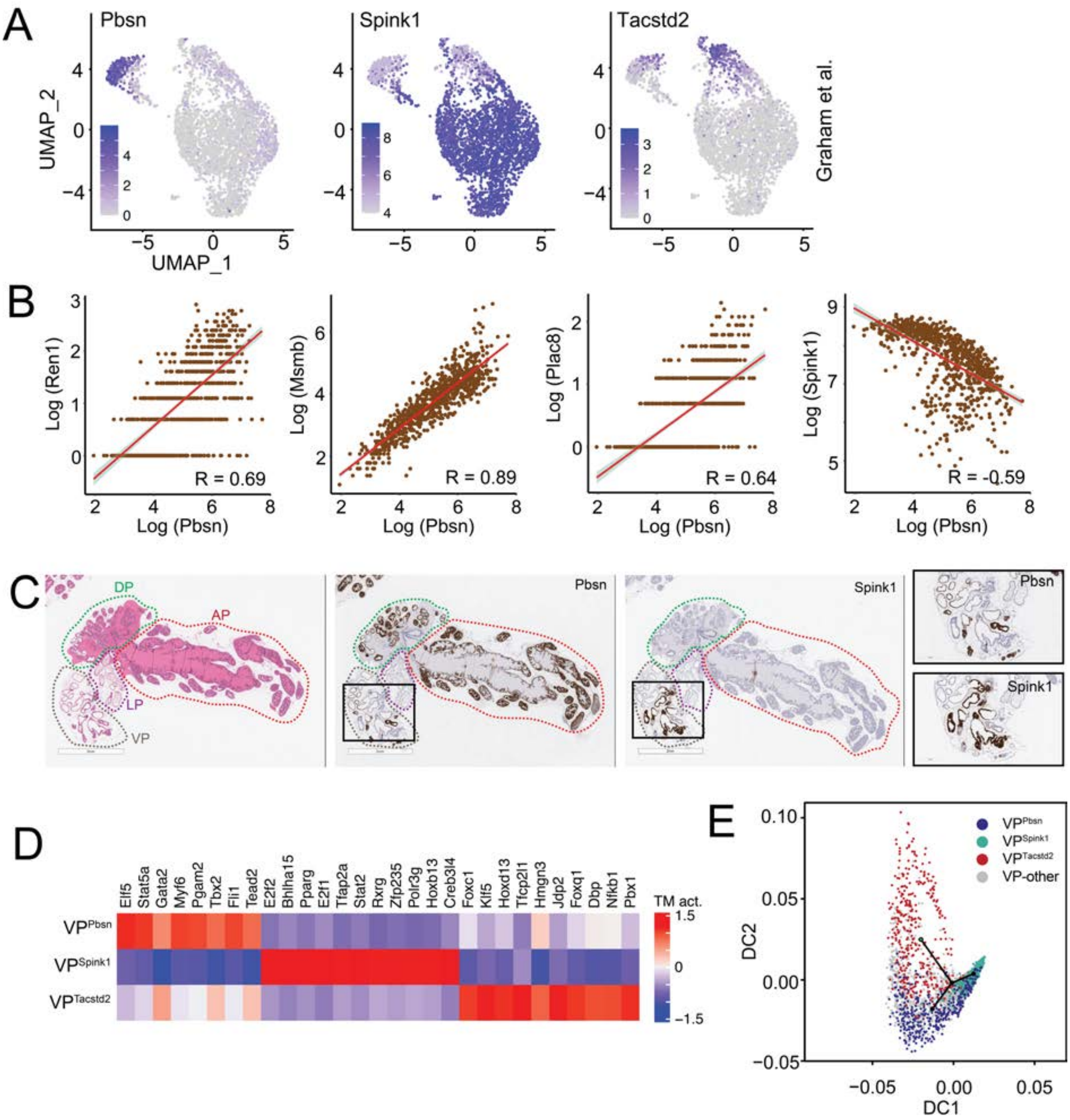


**Fig. S7. Related to figure 2. The VP has three distinct cell types, two enriched for distinct sets of androgen responsive genes and one enriched for stemness programs. (A)** UMAP of cells from the VP, as annotated in Graham et al. Left, *Pbsn*; middle, *Spink1*; and right, *Tacstd2* expression are represented as features (heatmap) on these UMAPs. **(B)** Correlation of *Pbsn* expression with *Ren1* (AP-specific), *Msmb* (LP-specific), *Plac8* (ADP-specific) and *Spink1* (VP-specific) expression in ST data. Correlation coefficients (R) are also depicted. **(C)** H&E image of whole mount mouse prostate with color coded anatomical locations annotated based on histological analysis. Representative single-color RNA-ISH image of a whole mount mouse prostate serial sections, with color coded outlines of lobes, wherein each section was stained for *Pbsn*, or *Spink1*. Scale bar, 2 mm. Zoomed-in region of specific glands in the VP (black outline) are also shown. Scale bar, 500 µm. **(D)** SCENIC-based heatmap of transcription module (TM) activity (TM act., columns) in ST spots from the VP (rows) that are enriched for *Pbsn* ($\mathrm{VP}^{\mathrm{Pbsn}}$), *Spink1* ($\mathrm{VP}^{\mathrm{Spink}}$) and *Tacstd2* ($\mathrm{VP}^{\mathrm{Tacstd2}}$) expression. **(E)** Diffusion plots from trajectory analysis of scRNAseq data of the VP from Graham et al, representing $Pbsn^{high}$, $Spink^{high}$ and $Tacstd2^{high}$ populations in the VP, i.e. $\mathrm{VP}^{\mathrm{Pbsn}}$ (blue), $\mathrm{VP}^{\mathrm{Spink1}}$ (cyan) and $\mathrm{VP}^{\mathrm{Tacstd2}}$ (red), respectively.

Green dot within the plot represents the start of the trajectory. Spots that could not be uniquely annotated as one of these transcriptional programs are in grey.

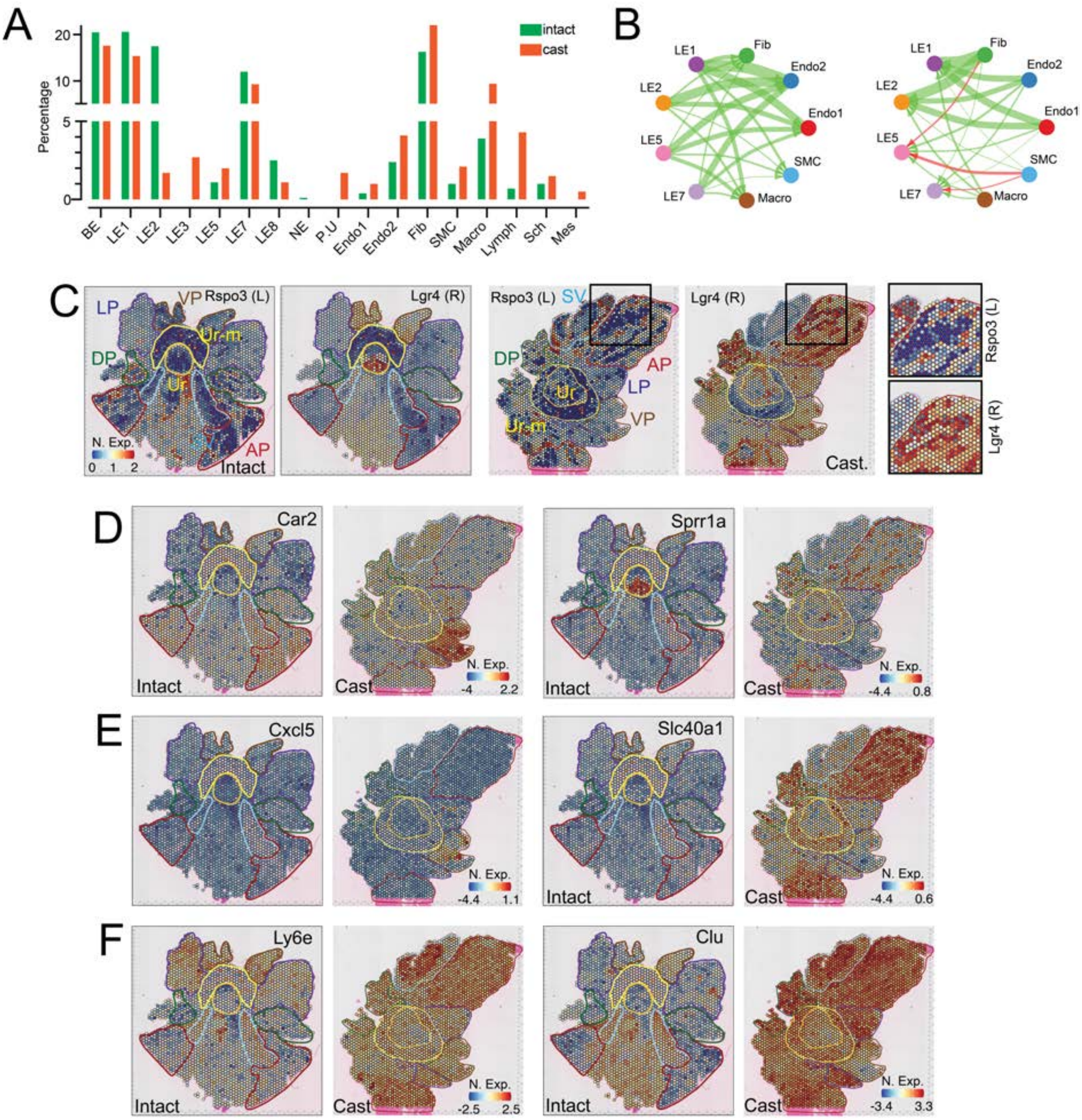


**Fig. S8. Related to figure 3. Castration drives dramatic reorganization of cell-specific transcriptomes. (A)** Bar plot depicting the percentage of each cell type represented in scRNAseq data from prostates of unperturbed (intact) and castrated (cast) mice. Cell types match those in Fig 3A. **(B)** Circos plots depicting ligand-receptor connectivity, based on scRNAseq data from intact and cast samples. Thickness of arrows depict the number of ligand-receptor connections with arrowheads depicting receptors. Left plot depicts ligand receptor interactions, with ligands on epithelia and receptors on the stroma and the right plot depicts vice-versa interactions. Green and red arrows depict connections gained and lost during castration respectively. **(C)** ST feature plots with color coded outlines of anatomical locations, representing the normalized expression (N. Exp.) of *Rspo3* (Ligand, L) and *Lgr4* (Receptor, R) in whole mount prostates from intact and cast samples. Zoomed-in region of the AP (black outline) from the cast sample is also shown. **(D-F)** ST feature plots with color coded outlines of anatomical locations, representing the normalized expression (N. Exp.) of *Car2* (ADP-specific in intact to VP-enriched in cast), *Sprr1a* (Ur-specific in intact to ADP-enriched in cast), *Cxcl5* (lowly expressed in intact to VP-enriched in cast), *Slc40a1* (lowly expressed in intact to ADP-enriched in cast), *Ly6e* (LVP-enriched in intact to ADLVP-enriched in cast) and *Clu* (lowly expressed in ADLVP, but enriched in non-prostatic regions of intact to ADLVP-enriched in cast) in whole mount prostates from intact and cast samples.

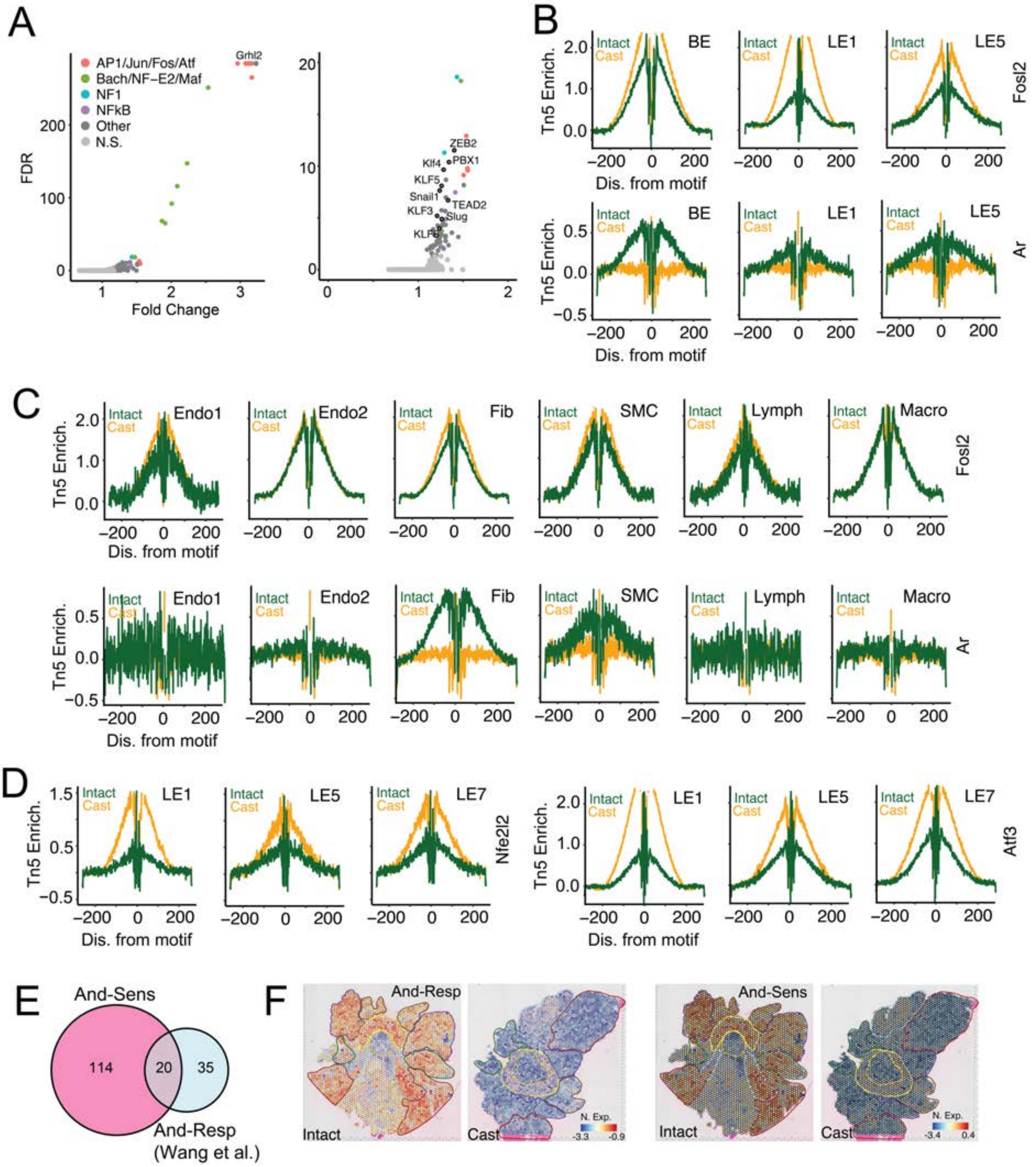


**Fig. S9. Related to figure 3. Castration induces stress responsive and stemness programs. (A)** Scatter plot of motif-associated accessibility changes (gains) versus false discovery rates (FDR) in LE cells from prostates of castrated mice, as compared to those from unperturbed mice. Each dot represents a motif and is color coded based on high-level grouping. Right, zoom-in of data on the left. **(B)** *Fosl2* (top) and *Ar* (bottom) footprints inferred from scMulti ATAC data (Tn5 Enrichment, Tn5 Enrich.) in other epithelial (BE, LE1 and LE5) cells from prostates of intact (green) and castrated (orange) mice. **(C)** *Fosl2* (top) and *Ar* (bottom) footprints inferred from scMulti ATAC data (Tn5 Enrichment, Tn5 Enrich.) in stromal (Endo1, Endo2, Fib, SMC, Lymph, Macro) cells from prostates of intact (green) and castrated (orange) mice. **(D)** *Nfe2l2* (left) and *Atf3* (right) footprints inferred from scMulti ATAC data (Tn5 Enrichment, Tn5 Enrich.) in LE (LE1, LE5, LE7) cells from prostates of intact (green) and castrated (orange) mice. **(E)** Venn diagram of androgen sensitive genes (And-Sens, identified by correlated reduction in LE cell gene expression and gene-proximal ARE accessibility changes during castration) and previously identified androgen response (And-Resp, Wang et al) gene set. **(F)** ST feature plots with color coded outlines of anatomical locations, representing the normalized expression (N. Exp.) of And-Resp and And-Sens gene sets in whole mount prostates from intact and cast samples.

Table S1. Meta-analysis comparing cell type annotations of the mouse prostate.

| This work (scRNAseq) | Crowley et al. 2020 | Guo et al. 2020 | Karthaus et al. 2020 | Joseph et al. 2020 | Graham et al. 2023 | Kirk et al 2024 | This work (ST) | This work (Top markers) |
|---|---|---|---|---|---|---|---|---|
| **LE1** | LumA | Luminal_B | Luminal_1 | AP | C57-AP, FVB-AP | L7, L8 | AP | *Ren1, C1rb, Glb1l3* |
| **LE2** | LumD | Luminal_B | Luminal_1 | AP | C57-DP, FVB-DP | L6 | AP, DP | *Ms4a5, Pate14, Tgm4* |
| **LE3** | | SV | SV_Luminal | SV/ED | | | SV, Ur-M, ADLVP, other | *Cyp4a12a, Crabp2, Pou3f3* |
| **LE4** | | | | | | | SV, Ur-M, ADLVP, other | *Svs1, Orm3, Dnase2b* |
| **LE5** | LumP | Luminal_C-Ur | Luminal_2Psca Luminal_3Foxi1 | Ur | | L5 | Ur + VP | *Psca, Krt4, Tacstd2* |
| **LE6** | | | | | | | SV, Ur-M, ADLVP, other | *Pip, Wfdc18, Bricd5* |
| **LE7** | LumV | Luminal_A | | VP | C57-VP, FVB-VP | L2, L3, L5 | VP | *Spink1, Sbp, Nupr1* |
| **LE8** | LumL | Luminal_A | | DLP | C57-LP, FVB-LP | L4 | LP | *Defb50 H2−Q10 Msmb* |
| **LE9** | | | | | | | SV, Ur-M, ADLVP, other | *Fxyd4, Acbd7, Cited1* |

Table S2. Meta-analysis comparing cell type annotations of prostates from castrated mice.

| This work (scRNAseq) | Kirk et al 2024 | This work (Top markers) |
|---|---|---|
| **C-LE1** | L1 | *Clu, Ly6e, Alcam* |
| **C-LE2** | | *Krt19, Slc40a1, Sprr1a* |
| **C-LE5** | L5 | *Psca, Krt4, Tacstd2* |
| **C-LE7, C-LE8** | L3 | *Car2, Piezo2, Cxcl5* |
